# Repurposing of Amisulpride, a known antipsychotic drug, to target synovial fibroblasts activation in arthritis

**DOI:** 10.1101/2022.08.02.500956

**Authors:** D. Papadopoulou, F. Roumelioti, C. Tzaferis, P. Chouvardas, A.K. Pedersen, F. Charalampous, E. Christodoulou-Vafeiadou, L. Ntari, N. Karagianni, M. Denis, J.V. Olsen, A.N. Matralis, G. Kollias

**Author notes:** Corresponding authors* Correspondence to George Kollias. Department for Biomedical Research, University of Bern, Bern, Switzerland (P. Chouvardas). BIOEMTECH, Lefkippos Attica Technology Park, NCSR “Demokritos”, Athens, Greece.

## Abstract

Synovial Fibroblasts (SFs) are key pathogenic drivers in arthritis and their *in vivo* activation by TNF is sufficient to orchestrate full arthritic pathogenesis in animal models. TNF blockade has been efficacious for a large percentage of Rheumatoid Arthritis (RA) patients, although characterized by a plethora of side effects. Novel therapeutic discoveries remain however challenging, especially in optimizing drug safety, side effects, longer-term responses, costs and administration routes. Aiming to find new potent therapeutics, we applied the L1000CDS^2^ search engine, in order to identify compounds that could potentially reverse the pathogenic expression signature of arthritogenic SFs, derived from the human TNF transgenic mouse model (*hTNFtg*). We identified a neuroleptic drug, namely Amisulpride, which was validated to reduce SFs’ inflammatory potential while decreasing the clinical score of *hTNFtg* polyarthritis. Notably, we found that Amisulpride did not exert its biological activities through its known targets Dopamine receptors 2 and 3 and Serotonin Receptor 7, nor through TNF-TNFRI binding inhibition. By applying a click chemistry approach, novel potential targets of Amisulpride were identified, which were further validated to repress *hTNFtg* SFs’ inflammatory potential *in vitro* (*Ascc3* and *Sec62*), while phosphoproteomics analysis revealed important fibroblast activation pathways, such as adhesion, to be altered upon treatment. Our data support that Amisulpride could provide an additive beneficial effect to patients suffering from RA and comorbid dysthymia, as it may reduce SFs pathogenicity in parallel with its anti-depressive activity. Importantly, Amisulpride may also serve as a “lead” compound for the development of novel, more potent therapeutics against chronic inflammatory diseases.

## 1. Introduction

Rheumatoid arthritis (RA) is a chronic inflammatory disease characterized by swelling and gradual destruction of the joints, that is marked by increased proliferation of resident mesenchymal cells (pannus formation) and expansion of inflammatory cells in the joint area, supported by increased angiogenesis*(1)*. Synovial fibroblasts (SFs) are one of the main cell types in the joints, which have been extensively associated with RA progression as they produce inflammatory cytokines/chemokines and degrading metalloproteinases, leading progressively to increased joint inflammation, stiffness and pain*(1)*.

One of the predominant cytokines that has been linked to RA pathogenesis is tumor necrosis factor alpha (TNF)*(2)*. We have shown that mice overexpressing TNF by either carrying a human TNF transgene (*hTNFtg* mice) or by deletion of ARE elements in the endogenous TNF gene (ΔARE mice)*(3)*, spontaneously develop chronic polyarthritis*(4)*, with histological manifestations fully resembling human RA. Importantly, TNF blockade could ameliorate disease onset and progression in both mouse models, while TNF signalling in SFs was found to be both necessary and sufficient for the orchestration of full pathology*(5, 6)*. Notably, *hTNFtg* pathogenic SFs were capable of initiating disease when transferred to the knees of healthy mice*(7)*, thus underlining the dominant and autonomous role of SFs in RA pathology initiation and progression. Notably, *hTNFtg* SFs, have been found to highly correlate with RA human Fibroblast Like Synoviocytes (FLS) not only in their gene expression profile*(8)* but also in their distinct roles in the joint area, with CD90^-^ SFs of synovial lining driving mainly bone destruction and CD90^+^ SFs of sublining area driving mainly inflammation*(9–11)*.

Current first line therapies against RA, including disease-modifying antirheumatic drugs (DMARDs), such as methotrexate, and targeted synthetic DMARDs inhibiting several kinases, such as Janus kinases (JAKs) or mitogen-activated protein kinase (MAPK), offer significant clinical benefits*(12)*. However, these therapeutics just remit the disease symptoms while the patients progressively lose response. In more severe cases, anti-TNF/IL6 biologics are prescribed, but their higher therapeutic potential is counterbalanced by decreased patient compliance due to invasive administration, as well as by a plethora of side effects, such as the gradual development of immunodeficiency and the production of antidrug antibodies, leading to further lower response to treatment *(13)*.

Herein, aiming to identify new potential therapeutics for RA treatment, we searched for compounds in a repurposing setting, as this may provide candidates that have been already assessed for their toxicity, formulation and the route of administration*(14)*. The identification of repositioned compounds has now become less dependent on serendipitous observations, since a multitude of computational and experimental approaches have been developed.

To this end, the L1000CDS^2^ search engine*(15)*, employed successfully in the past for repurposing small molecules as therapeutics for several diseases*(16)*, was used here in order to identify compounds that could potentially reverse the pathogenic gene expression signature of *hTNFtg* SFs. Using this methodology, a commercially available neuroleptic medicine, namely Amisulpride (Solian, Sanofi), was identified and validated further to downregulate *hTNFtg* SFs’ activation by reducing both their inflammatory and adhesive potential. Interestingly, the mechanism through which Amisulpride exerts its activity was found to be independent of Dopamine receptor-2 and -3 (DRD2 and DRD3) and Serotonin receptor-7 (HTR7) function, the main reported targets of the drug. By applying a click chemistry approach, five novel potential targets of Amisulpride were identified, two of which, namely *Ascc3* and *Sec62*, were further validated to repress *hTNFtg* SFs’ inflammatory potential *in vitro*. Notably, *in vivo* experiments using the *hTNFtg* mouse model demonstrated the amelioration of polyarthritis by Amisulpride. Collectively, treatment with Amisulpride may offer an added advantage to the treatment of RA by addressing simultaneously RA comorbid depression. Additionally, Amisulpride may serve as a “lead” compound for the development of novel, more potent and selective therapeutics against chronic inflammatory disorders.

## 2. Results

### 2.1 Identification of Amisulpride as a modifier of hTNFtg SFs activation

To identify new drug candidates as potential therapeutics for RA, the publicly available search engine “L1000CDS^2^”, which explores a set of different combinations of treatments and cell lines, aiming to find perturbations that either mimic or reverse user’s input signature, was used *(15)*. As shown in Figure 1A (summarized strategy followed), initially 3’ mRNA sequencing of SFs isolated from 8-week old *hTNFtg* (established disease) and wild type (WT) controls was performed and the differentially expressed genes were identified. The results of this analysis were then combined with data already published*(8)*, aiming at precisely determining those genes that are commonly deregulated at all 3 stages of the disease (early, established, late) when compared with the WT control (Figure S1A). Additionally, the expression profile of 8-week *hTNFtg* SFs after a 48h treatment with 1μg/ml Infliximab, a widely used anti-TNF biologic, was compared with the relevant untreated *hTNFtg* SFs (Figure S1B). Only one drug, namely Amisulpride, was predicted to commonly reverse the disease signature and mimic the Infliximab effect on arthritogenic SFs (Figure S1C). Figure S1D presents the fifty top candidate perturbations that were proposed by “L1000CDS^2^”*(15)* search engine, respectively. Thus, Amisulpride was examined as the most promising candidate compound in terms of targeting genes activated upon chronic inflammation.

**Figure 1:**
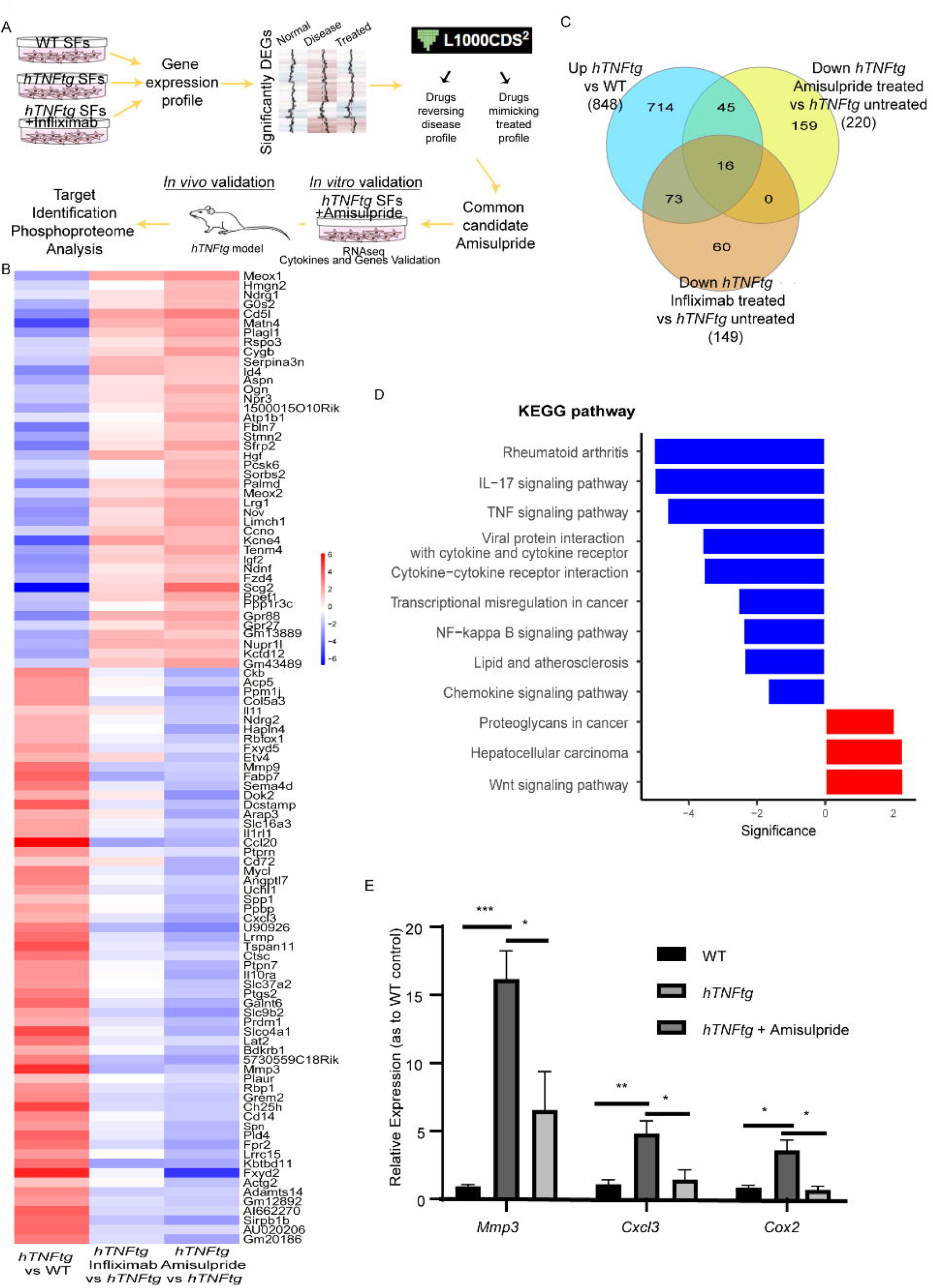
Amisulpride treatment affects the TNF pathway in *hTNFtg* SFs. A) Pipeline used to identify and test Amisulpride as modifier of *hTNFtg* SFs’ activation. B) Heatmap of the 103 genes that are deregulated in *hTNFtg* SFs vs WT and are restored upon Amisulpride treatment. C) Venn diagram showing the genes being upregulated in *hTNFtg* SFs vs WT while being downregulated upon Amisulpride/ Infiliximab treatment. D)Kegg pathway analysis of the 103 genes described in B. E) Validation of RNAseq identified genes through quantitative Real Time PCR. (* p-value < 0.05; ** p-value < 0.01; *** p-value ≤ 0.0001, all data are shown as mean ± SEM and all comparisons were made against *hTNFtg* vehicle sample using Student’s t test).

### 2.2 Validation of Amisulpride as a modifier of hTNFtg SFs activation

Amisulpride is an antipsychotic drug acting as a powerful antagonist of DRD2/ DRD3 and HTR7, being administered in high doses to treat schizophrenia and in low doses to treat depressive disorders*(17)*.

We further validated the effect of Amisulpride in *hTNFtg* SFs by comparing the expression profile of *hTNFtg* SFs treated with Amisulpride for 48h to that of untreated *hTNFtg* SFs. Amisulpride restored a significant number of genes *hTNFtg* SFs (103 genes in total that are deregulated in hTNFtg SFs vs WT controls) including downregulation of genes expressing several inflammatory chemokines and Metalloproteinases (MMPs) (Figure 1B and 1C). Notably, KEGG pathway analysis of these genes, revealed RA pathogenesis and TNF signalling pathway among the top enriched functional terms (Figure 1D). The expression of genes such as *Mmp3, Cxcl3* and *Cox2*, that are important in joint inflammation and bone destruction was also validated by quantitative real time PCR (Figure 1E) and it was found to be decreased following Amisulpride treatment of arthritic SFs. Importantly, genes that were restored following treatment with Amisulpride, were also commonly regulated by Infliximab treatment on *hTNFtg* SFs, validating further our identification procedure (Figure 1B and 1C).

Notably, Amisulpride treatment resulted in overexpression of 42 genes that are found to be downregulated in *hTNFtg* vs WT SFs (Figure 1B). Analysis of these genes highlighted Wnt signalling as the top enriched pathway which interestingly, has been also found to be deregulated in human RA and in relevant experimental animal models of the disease*(18, 19)*, while it has been suggested to have a protective role in *hTNFtg* mouse model*(20)*.

The anti-inflammatory profile of Amisulpride was further confirmed at the protein level by the dose-dependent reduction of the elevated pro-inflammatory chemokines CCL20/ MIP3 and CCL5/ RANTES in the supernatants of *hTNFtg* SFs, upon drug treatment (Figure 2A). A similar effect of Amisulpride on CCL5 and CCL20 production was also noticed in WT SFs exogenously stimulated by TNF (Figure 2B), indicating that Amisulpride interferes with TNF-driven cell responses both at a chronic and at an acute TNF stimulation setting. Of note, all drug concentrations used were non-toxic according to a crystal violet assay performed (Figure S2A). Moreover, the levels of both the monocyte chemoattractant protein-1 (MCP1), CCL2, and the angiogenic chemokine CXCL5 were found to be reduced in the supernatants of treated *hTNFtg* SFs, indicating that Amisulpride has a wide spectrum of anti-inflammatory properties (Figure S2B). Amisulpride also reduced both the transcription and the secretion of hTNF by *hTNFtg* SFs, suggesting that the drug affects *in vitro* TNF production which drives *hTNFtg* SFs pathogenicity (Figure 2C). Finally, Amisulpride was able to effectively inhibit the TNF-induced cytotoxicity of L929 cells*(21)* (Figure 2D), providing further evidence on the potential implication of the drug in the TNF signalling pathway, rendering it as a promising candidate for further *in vivo* use in the *hTNFtg* polyarthritis model.

**Figure 2:**
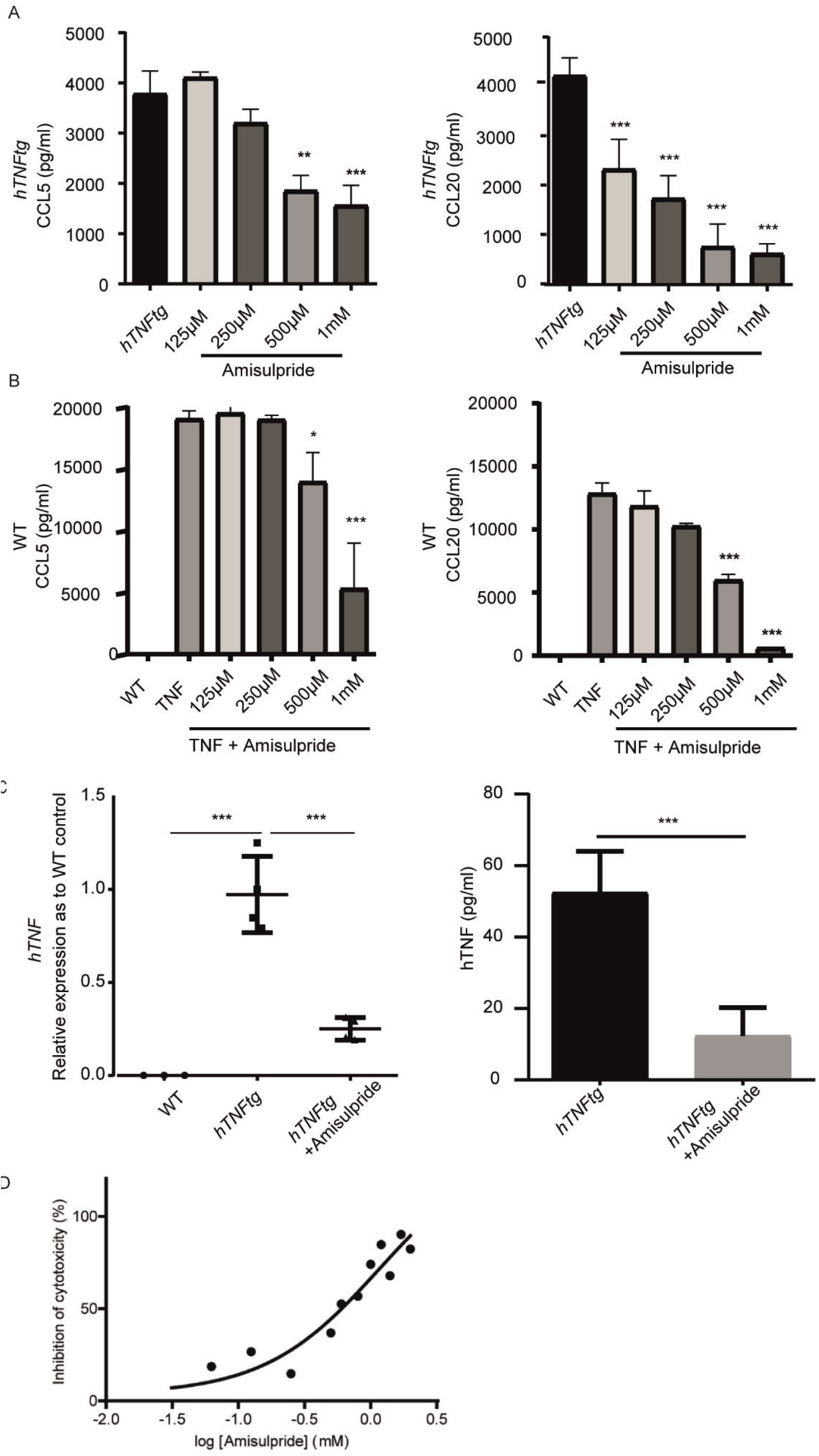
The anti-TNF activity of Amisulpride. Chemokines detection in supernatants of A) *hTNFtg* SFs and B) WT SFs stimulated with 10ng/ml hTNF, treated with Amisulpride at the indicated concentrations for 48h. C) Amisulpride affects both the transcription and the secretion of hTNF in *hTNFtg* treated SFs. D) L929 cells TNF induced necroptosis is inhibited by Amisulpride in a dose dependent manner. (* p-value < 0.05; ** p-value < 0.01; *** p-value ≤ 0.0001, all data are shown as mean ± SEM and all comparisons were made against *hTNFtg* vehicle sample using Student’s t test.)

### 2.3 Amisulpride alleviates acute and chronic inflammation in vivo

Amisulpride known targets are DRD2/ DRD3 and HTR7, which although they are best known as neurotransmitters, they have been implicated in immune system function, inhibiting acute inflammation in vivo*(22, 23)*.

Similarly, the anti-inflammatory activity of Amisulpride *in vivo* was validated in the LPS acute sepsis experimental model in wild type mice, where the administration of Amisulpride downregulated significantly in a dose-dependent manner the elevated serum levels of mouse TNF and IL6 detected 1.5h post induction (Figure 3A).

**Figure 3:**
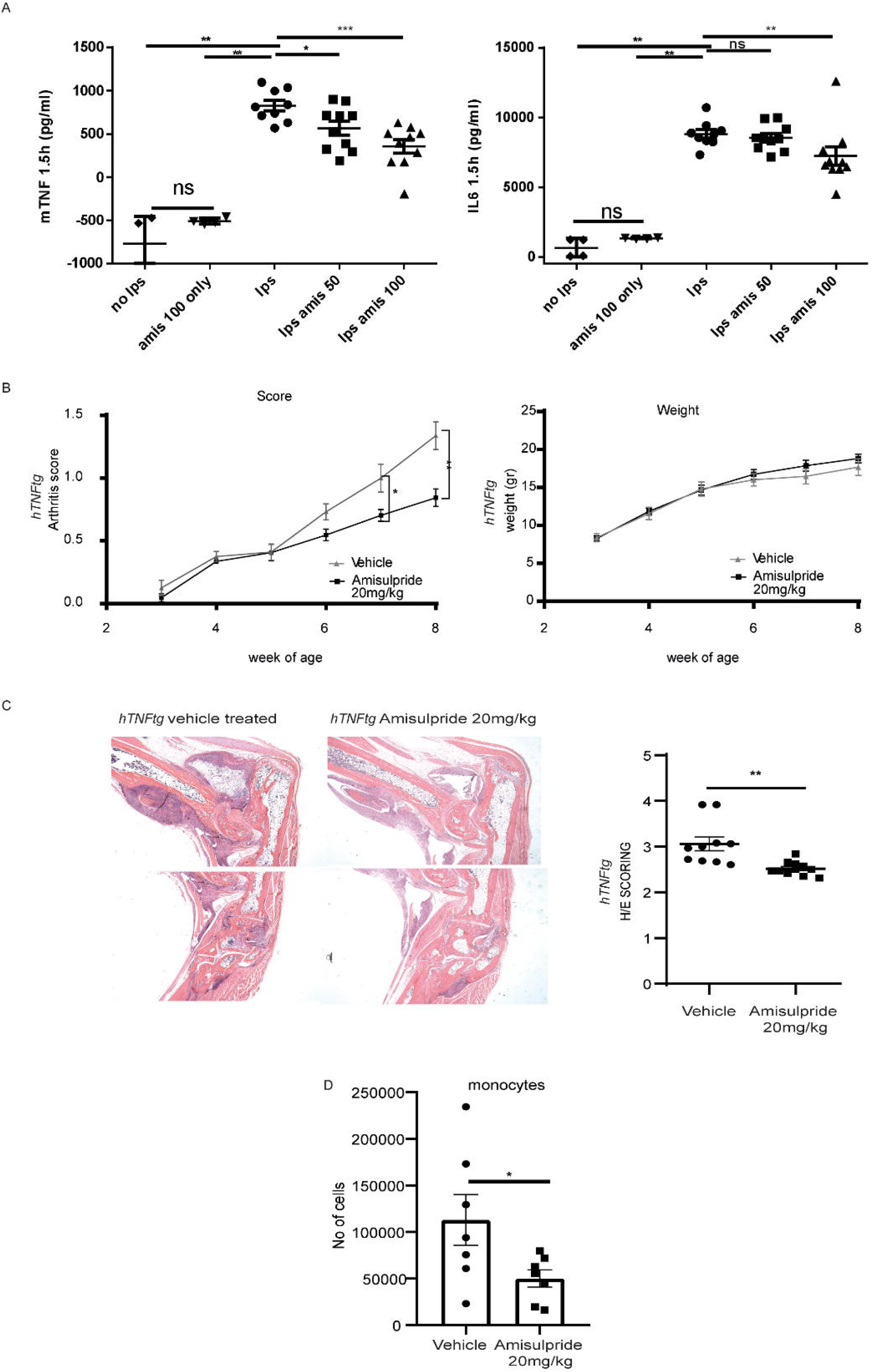
In vivo effect of Amisulpride. A. Amisulpride treatment in LPS induced acute sepsis model significantly downregulates elevated serum mTNF and IL6, when compared with the vehicle treated controls. 5weeks of Amisulpride treatment in *hTNFtg* model significantly downregulates B. arthritis clinical score C. synovitis scoring of joints’ in H/E stained paraffin sections and D. monocytes numbers in FACs-based immune infiltration analysis, when compared with vehicle treated controls. (* p-value < 0.05; ** p-value < 0.01; *** p-value ≤ 0.0001, all data are shown as mean ± SEM and all comparisons were made against *hTNFtg* vehicle sample using Student’s t test.)

Considering the inhibitory activity of Amisulpride on the TNF signalling pathway, both in SFs and in the LPS-induced acute inflammation, the therapeutic potential of the drug was further evaluated *in vivo* in the *hTNFtg* erosive polyarthritis *(24)*, which has been previously used successfully for drug testing*(25)*. Amisulpride was administered orally twice per day at a prophylactic mode (before the onset of the disease until the established disease state). The selected dose of Amisulpride was based on the known toxicity data of the drug and the relevant human dose prescribed for its original use in the field of depressive disorders*(17, 26)*. Amisulpride decreased the clinical arthritis score of *hTNFtg* mice, decreasing also significantly the synovitis score in H/E-stained histological sections (Figure 3B and 3C). Intriguingly, the significant anti-arthritic activity of Amisulpride was not associated with a significant effect in either the bone erosion or the cartilage destruction of the treated animals (Figure S3A and S3B), indicating the potential involvement of the drug in attenuating mainly the influx of inflammatory cells in the joints of arthritic mice. Weekly body weight assessment and gross behavioural effect observation confirmed the already described safety profile of the drug (Figure 3B).

FACs based quantification of infiltrated CD45^+^ cells, macrophages, monocytes, neutrophils, eosinophils, dendritic cells, CD4^+^ T cells and CD8^+^ T cells in the ankle joints of 8 week-old mice treated with Amisulpride showed that the drug attenuates mainly the numbers of monocytes when compared with vehicle treated controls (Figure 3D and S3C), which can be further related to the decreased MCP1 levels detected on the hTNFtg SFs when treated with the drug.

Importantly, we did not find any significant difference in the hTNF serum levels of treated and untreated mice (Figure S3D), indicating that *in vivo* other factors seem to control the improved clinical outcome observed in *hTNFtg* mice.

### 2.4 Amisulpride effect on joint fibroblasts is mediated neither through its known targets DRD2, DRD3 and HTR7, nor through TNF-TNFRI binding inhibition

We next investigated the molecular mechanism of Amisulpride function. We first assessed whether Amisulpride effect is mediated through its known targets DRD2/DRD3/HTR7 receptors. Although B- and T-lymphocytes, macrophages and dendritic cells have been shown to express dopamine receptors*(27)*, the expression of these receptors in RA FLS remains controversial*(28, 29)*. Interestingly, in a recent study their expression seemed to affect mainly the RA FLS’ migration while their expression has been associated with the patients ‘age*(30)*. Importantly, in our study *Drd2*/*Drd3*/*Htr7* genes were not expressed neither in *hTNFtg* cultured SFs nor in fresh *hTNFtg* isolated fibroblasts, of both CD90^-^ or CD90^+^ subset*(9)* (Figure 4A). Thus, it seems that Amisulpride targets other than its main receptors on *hTNFtg* SFs.

**Figure 4:**
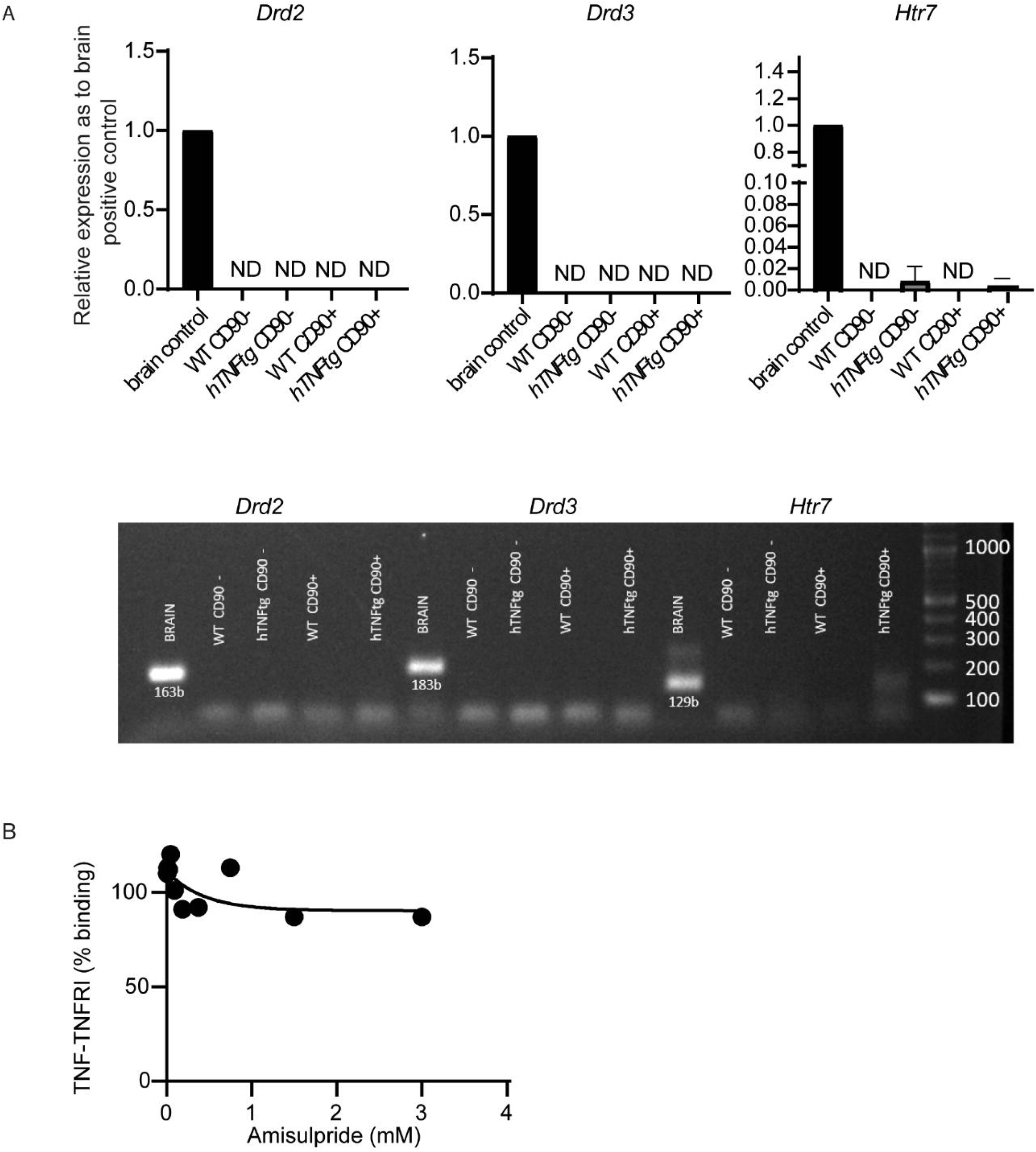
Amisulpride effect on joint SFs is neither mediated by its known targets nor through TNF-TNFRI binding inhibition. A. *Drd2*/ *Drd3*/ *Htr7* are not expressed on CD31-/ CD45-/PDPN+/ CD90- or CD90+ *hTNFtg* fresh isolated SFs. ND: Not Detected. Brain sample was used as a positive control. B. TNF-TNFRI binding upon Amisulpride.

Additionally, and since TNFRI expression on SFs has been found crucial for *hTNFtg* pathology*(6, 31)*, the ability of the drug to directly interrupt the interaction of TNF with its receptor TNFRI was quantified. Nevertheless, as Figure 4B depicts, Amisulpride does not suppress the binding of TNF to its main receptor TNFR1, indicating that the anti-TNF effect of the drug is not mediated through the TNF-TNFR1 binding inhibition.

Conclusively, our results suggest that Amisulpride activity on *hTNFtg* SFs is not mediated by the main known targets of the drug (DRD2, DRD3 and HTR7), while its effect on TNF signalling is not exhibited through the interruption of binding of TNF to TNFRI.

### 2.5 Chemoproteomic identification of potential molecular targets of Amisulpride

Our efforts were then directed towards setting up a target identification protocol using the pathogenic *hTNFtg* SFs as a cellular context.

We first designed and synthesized two bioactive chemical probes which structurally resemble Amisulpride (click compounds), while containing a click handle in a different position (triple bond in Figure 5A). The click compounds can ligate to biotin and pull down the potential target proteins after being bound to streptavidin beads, thus enabling target identification through LC-MS/MS analysis*(32, 33)*.

**Figure 5:**
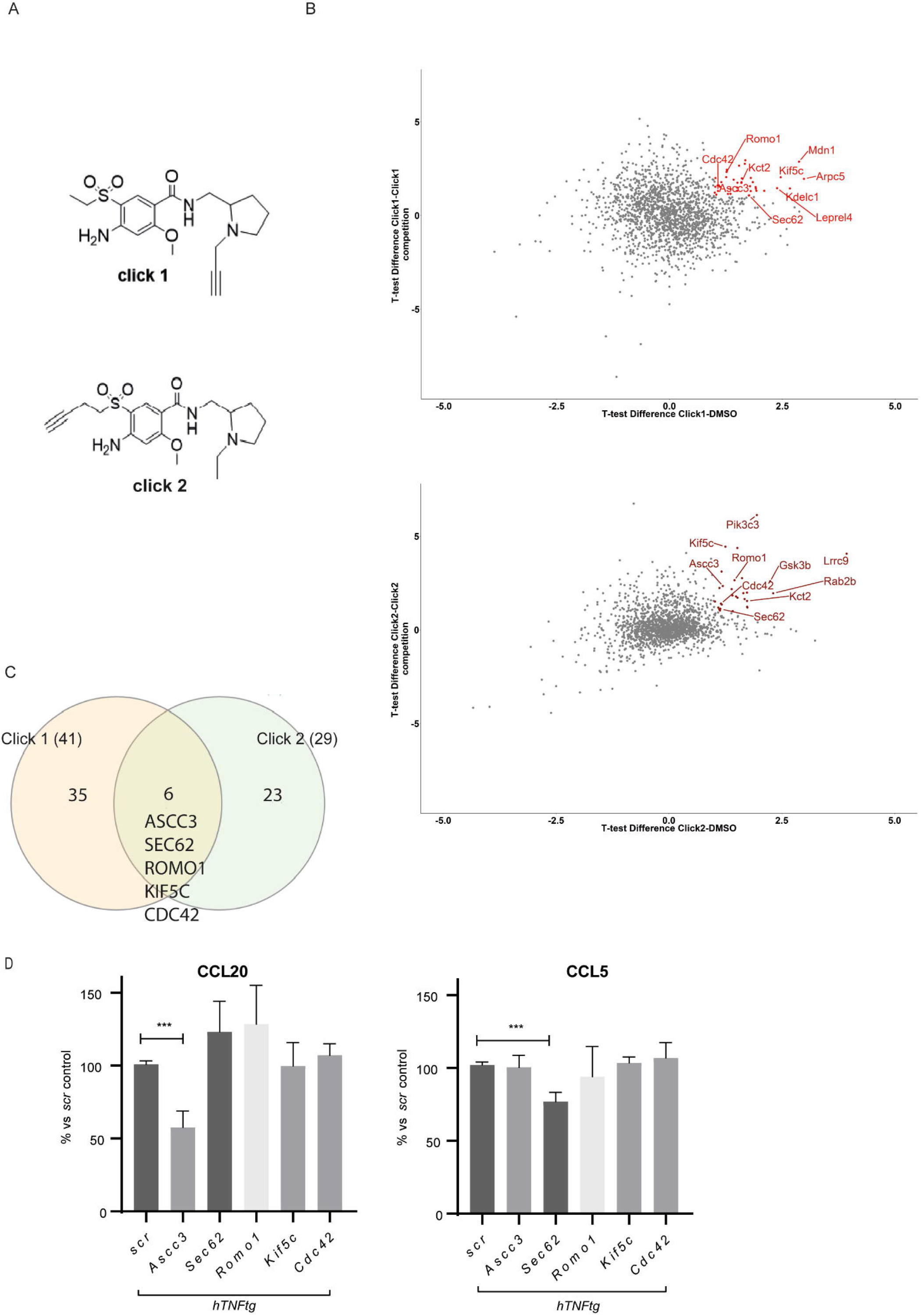
Drug targets identification. A Structure of the two synthesized click compounds B Volcano plots with highlighted candidates, combining differences (Student’s t test difference>1) in proteins identified in each active probe sample but not in the DMSO or the competition control. C Venn diagram of the proteins identified in B. D. ShRNAs mediated deletion of *Ascc3, Sec62, Romo2, Kif5c* and *Cdc42*, identified that combined targeting of ASCC3 and SEC62 support the anti-inflammatory profile of the drug. (* p-value < 0.05; ** p-value < 0.01; *** p-value ≤ 0.0001, all data are shown as mean ± SEM and all comparisons were made against Scramble-scr sample using Student’s t test.)

We synthesized two click probes in order to exclude as many artefacts as possible and to end up with a low number of potential protein targets for validation. In the first click probe (click 1, Figure 5A), the alkyne was incorporated into Amisulpride as substituent of the pyrrolidine group, while in the second as substituent of the sulfone group (click 2, Figure 5A). The optimized synthetic route followed for the synthesis of the two click probes is displayed in Figure S4A and S4B. Figure S5 includes the characterization of the synthesized Amisulpride and its click derivatives (click 1 and click 2) by ^1^H-NMR and mass spectometry.

In order to confirm that both click compounds retain their bioactivity we repeated the CCL5 and CCL20 quantification in *hTNFtg* SFs treated with each individual probe. Both click analogues exhibited a similar to or even better (Figure S4C) than Amisulpride effect (Figure 2A).

Hence, the *hTNFtg* SFs were treated with each click derivative (25 μM), while two negative controls were used. The first control included *hTNFtg* SFs treated with vehicle (Dmso, no probe). The second control (Competition) included *hTNFtg* SFs treated with each click derivative (25 μM) plus an excess concentration (10x, 250μM) of Amisulpride, which would compete off each probe (click 1/ click 2) and potentially occupy the target binding sites. The target proteins were identified by mass spectrometry after being conjugated with a biotin tag and pulled down on streptavidin beads. The mass spectrometry results were further analysed by a) plotting the protein enrichment by each probe (click 1/ click 2) compared to vehicle (dmso) and b) plotting the proteins enriched by click 1/ click 2 compared to the competitor (Amisulpride) (Figure 5B). Combining differences (Student’s T test difference >1) in proteins identified in both active probe samples but not in the two negative controls (dmso/ Amisulpride) delivered a shortlist of potential targets (Figure 5C, Venn diagram). The six common candidates identified were ASCC3, SEC62, ROMO1, KIF5C, CDC42 and KCT2 (Figure 5C). KCT2 was then excluded from further analysis, as it is a protein expressed mainly by keratinocytes, indicating that it is probably a contaminant protein in the experiment.

In conclusion, we found that Amisulpride possibly regulates the activation of *hTNFtg* SFs by binding to ASCC3, SEC62, ROMO1, KIF5C and/ or CDC42.

### 2.6 Validation of potential molecular targets of Amisulpride

The identified targets seemed potentially capable off ameliorating *hTNFtg* SFs’ pathogenicity. Activating signal co-integrator 1 complex subunit 3 (*Ascc3*) encodes a 3′-5′ DNA helicase, while in cell lines loss of *Ascc3* leads to reduced cell proliferation*(34)*. Kinesin heavy chain isoform 5C (KIF5C) is located in microtubules, where among other proteins, it has been found necessary for the transport of N-cadherin between the Golgi and the plasma membrane, facilitating cell to cell adhesion in fibroblasts*(35)*. Cell division cycle 42 (CDC42) has been also associated with adherence of fibroblasts as it belongs to a subfamily of Rho GTPases*(36)*, while it has been previously linked to antidepressive compounds, that act through targeting of the HTR7 receptor*(37)*. ROS modulator 1 (ROMO1) is localized in the mitochondria, being responsible for TNF*a*-induced ROS production*(38)*. Lastly, SEC62 is located in the endoplasmic reticulum (ER) and is responsible, as a member of SEC61/62/63 complex, for proteins translocation in the ER and for cell calcium regulation*(39)*.

To verify further the role of the 5 proteins identified as potential molecular targets of Amisulpride in the *hTNFtg* arthritic SFs, we designed shRNAs to delete each one of their corresponding genes using the Lenti-X Lentiviral expression system. After confirming the downregulation of each mRNA (Figure S4D), we checked which of the potential targets could successfully downregulate both CCL5 and CCL20, resulting in an anti-inflammatory effect similar to Amisulpride (Figure 5D). Of note, deletion of *Sec62* and *Ascc3* significantly reduced pathogenic chemokines levels (CCL5 and CCL20, respectively). This result indicates that the amelioration of *hTNFtg* SFs inflammatory activation offered by Amisulpride is through targeting of both ASCC3 and SEC62.

The anti-inflammatory role of ASCC3 and SEC62 targeting can also be correlated with the effect of Amisulpride on *hTNFtg* SFs gene signature (Figure 1D). Analysis of expression profile of treated *hTNFtg* SFs revealed TNF/ NF-κB pathway and chemokine signalling as main cascades regulated by the drug, confirming that the molecular targets of the drug may play a role in response to inflammatory signals.

In conclusion, five novel potential Amisulpride targets on *hTNFtg* SFs were identified and we propose herein that the anti-inflammatory profile of the drug is supported by the dual targeting of Assc3 and Sec62.

### 2.7 Amisulpride influences pathways known to be implicated in arthritogenic activation of SFs

To gain a deeper insight into the cellular signaling affected by the drug, phosphoproteomic profiling of treated synovial fibroblasts was deployed*(40)*. hTNF-induced WT SFs rather than *hTNFtg* SFs were used in order for the cells to synchronize in the context of TNF stimulation and signaling. Three different time points of hTNF induction were used and compared to cells pretreated with Amisulpride 1h before being stimulated with hTNF at similar time points (5’, 15’, 30’).

Initially, all treated samples were clustered together and were compared versus the untreated ones. Figure S6A displays a volcano plot that integrates the deregulated phosphosites in treated samples when compared with the respective untreated controls. Notably, hierarchical clustering revealed clearly distinguishable groups of phospho-regulations that were dependent on the presence of the inhibitor (Figure 6A).

**Figure 6:**
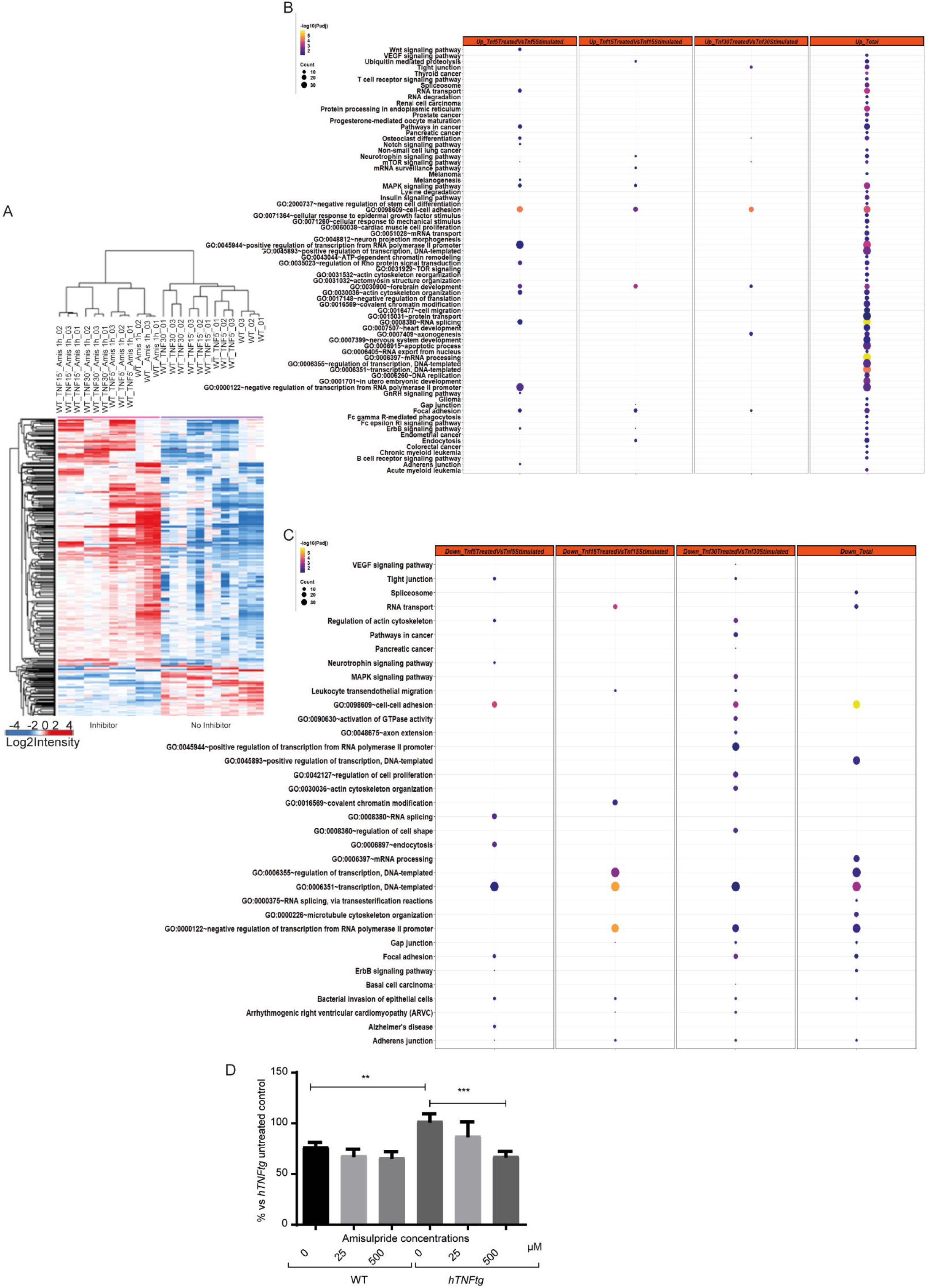
Phosphoproteomics analysis of intraarticular SFs treated with Amisulrpide reveal pathways that are regulated upon the drug treatment. A. Heatmap based on hierarchical clustering analysis of t-test significantly regulated phosphosites comparing Amisulpride treated and untreated SFs reveals clearly distinguishable row-clusters depending on inhibitor-effect. Kegg pathways and GO terms enrichment analysis for B. upregulated and C. downregulated phosphorylations upon Amisulpride treatment. D. Amisulpride downregulates the adherence of activated *hTNFtg* SFs. (* p-value < 0.05; ** p-value < 0.01; *** p-value ≤ 0.0001, all data are shown as mean ± SEM and all comparisons were made against *hTNFtg* vehicle treated sample)

Moreover, the phosphosites affected after treatment of WT fibroblasts with 10 ng/mL hTNF were identified in three different time points. These phospho-regulations were subsequently compared with the respective ones after pre-treatment of the cells with Amisulpride (500 μM) 1h before each relevant stimulation (volcano plots at Figure S6B and SC). Both the Kegg pathways and the Gene ontology terms enriched in all comparisons were analysed (Figure 6B and 6C).

Most of the phosphosites were upregulated upon inhibitor treatment, highlighting cell adherence, focal adhesion and MAPK signalling as implicated processes, pathways that are known to play a role in pathogenic fibroblasts’ activation (Figure 6B). Focusing on the downregulated phospho-changes, similar processes were found to be involved, containing cell to cell adhesion, focal adhesion and adherence junction as well as regulation of transcription (Figure 6C).

Notably, adhesion is known to be enhanced in pathogenic *hTNFtg* fibroblasts when compared to WT controls*(41)* and Amisulpride was found to significantly reduce cell adherence, therefore confirming the pathways proposed to be regulated by the drug in the phosphoproteome analysis (Figure 6D).

Interestingly, some of the identified protein targets of the drug (Figure 5C) can be highly connected with the pathways regulated upon Amisulpride treatment on SFs. Specifically, the phosphorylation of SEC62 (at Ser341), validated as one of Amisulpride targets on SFs, seems to be upregulated upon treatment, indicating that probably the endoplasmic reticulum translocation pathway is affected by the drug (Figure 6B). In addition, another potential drug target, CDC42, is a Rho GTPase, known to regulate filopodia formation and adherence of fibroblasts*(36, 42)*. Importantly, Rho protein signal transduction, cell to cell adhesion and actin cytoskeleton organisation were found to be upregulated too by Amisulpride, as seen in Figure 6B. Moreover, CDC42EP1 (MSE55), a CDC42 binding protein, that regulates cytoskeleton dynamics, is dephosphorylated (at S371 and S207) upon Amisulpride treatment, underlining further cell to cell adhesion alteration upon treatment (Figure 6C). Finally, ASCC3, as an RNA helicase, is implicated in transcription regulation, one of the top pathways identified in the phospho-proteome analysis (Figure 6C).

On the whole, *hTNFtg in vivo* polyarthritis amelioration by Amisulpride seems to be mediated by the targeting of the drug against *hTNFtg* SFs’ inflammatory and adhesive signalling, in which the aforementioned identified novel molecular targets appear to play an important role.

## 3. Discussion

In this study we have identified Amisulpride, a known antipsychotic agent known to inhibit dopamine and serotonin receptors, as a modifier of arthritic fibroblasts’ activation and mouse polyarthritis disease. RA is characterized by joints’ sensory nerve fibers accumulation that can respond to antidepressants agents*(43)*. Although neurotransmitters are considered as key role molecules in signal transduction of nervous system, they have been also implicated in immune cells’ regulation*(44)*. In this context, anti-dopamine drugs have been used to supress sepsis*(45)*, stress-induced neuroinflammation*(46)* and pro-inflammatory cytokines production in lipopolysaccharide (LPS)-induced macrophages*(47)*, while they have exhibited differential results in several RA *in vivo* studies*(48)*.

Amisulpride is widely used in high doses to treat schizophrenia and in low doses for treating depressive disorders*(49, 50)*. In the present study, it has been demonstrated that low doses of the drug exert, additionally, an anti-rheumatic potential. The new activity of the drug *in vivo* could be synergistic. It could first be ascribed to DRD2, DRD3 and HTR7 antagonism (Amisulpride known targets) on joint cells that express these receptors and their consequent response to dopamine expressed by joints SFs*(51)*. Furthermore, it could be due to the modification of function of the five newly identified drug off-targets on arthritogenic fibroblasts, two of which (ASCC3 and SEC62) have been validated here to support drug’s anti-inflammatory effect. This dual role underlines the high potential of this specific drug to be repurposed. Considering that the *de novo* identification of RA therapeutics is both costly and time consuming, drug repositioning consists a tractable alternative, since essential issues such as bioavailability, toxicity and manufacturing routes of the already tested drugs are known and established*(14)*.

Regarding the new drug targets validated to underlie the anti-inflammatory function of Amisulpride, ASCC3 is the largest subunit of ASC1 complex, where it functions as a 3′-5′ DNA helicase, together with ASCC1, ASCC2 and ASC1/ TRIP4*(34, 52)*. Inhibitors targeting DNA/ RNA helicases have been proposed to regulate viral, bacterial or cancer cells’ proliferation and responses*(80, 81)*. Ascc3 is also considered to participate in DNA repair pathways, as it prepares single-stranded DNA for AlkBH3 to proceed in de-alkylation repair*(34)* and it resolves stalled ribosomes*(55, 56)*. Moreover, ASC1 complex has been involved in transcriptional regulation, regulating alone or through transcription integrators SRC-1 and CBP-p300, several transcription factors such as CREB, STATs, AP-1 and NF-κB*(52, 57)*. ASCC3 involvement in the regulation of these inflammation-related transcription factors*(58)* can be further substantiated by the findings of the present study, where *Ascc3* downregulation is able to cause a significant reduction of the pro-inflammatory CCL20 secretion in TNF-induced activated SFs.

The translocation protein SEC62, on the other hand, is part of the dimeric SEC62/SEC63 complex, that along with Sec61 is located in the membrane of the endoplasmic reticulum (ER), facilitating the translocation of nascent polypeptides into the ER and the cell calcium homeostasis*(39, 59, 60)*. Additionally, SEC62 has been reported to play a crucial role in the successful ER-stress response, as it favours the release of the accumulated ER chaperones driving cell physiological homeostasis*(60)*. Notably, SEC62 has been reported as oncogenic, being overexpressed in a variety of tumours, supporting the migration but not the proliferation of several cancer cell lines*(61)*. The supportive effect of SEC62 in tumour metastasis can be attributed either to the deregulated translocation of migration-related precursor proteins at the ER or to the inhibition of Ca^2+^ homeostasis along with the attributed ER stress tolerance*(39)*. In particular, no SEC62 specific inhibitors have been reported to date, but inhibition of Sec62 function through antagonizing cellular Ca^2+^ homeostasis by CaM antagonists has been proven to mimic *SEC62* deletion *in vitro* by inhibiting migration and proliferation of human tumour cells*(61, 62)*. Targeting of SEC62 could also be beneficial in RA, since SEC62 has been found to be induced in a human RA synovial tissue study, and especially in a patients’ cohort that is characterized by expansion of FLS and not by predominance of myeloid/lymphoid compartment before receiving any treatment*(63)*. This subgroup of patients can be probably simulated by *hTNFtg* mice where TNF signalling in fibroblasts specifically is sufficient and necessary to initiate disease*(5, 6)*.

Psychiatric disorders are common comorbidities of RA and almost 19% of RA patients develop depression, a percentage much higher than the one met in the general population*(64)*. This comorbidity affects not only patients’ daily life but also the physical aspects of the disease as well as response to therapies*(65, 66)*. A common association of RA with its comorbid depression could be the fatigue and the pain that patients suffering from chronic inflammation can face*(67, 68)*. Although this might be the case, it has also been proven that the immune system itself can mediate depression pathogenesis, as circulating cytokines, such as IL6 and TNF, can activate endothelial cells of blood brain barrier, thus enabling the circulating mediators to enter in the central nervous system and to cause several mental disabilitites*(69)*. Additionally, pain and fatigue themselves can activate immune response which can in turn cause dysthymia pathology*(70, 71)*. Of note, anti-TNF biologics have been proposed for alleviating both RA and depression symptoms*(68)*. However, biologics prescription is approved only in severe cases of RA and not all patients respond to them, thereby necessitating the frequent prescription of additional antidepressants*(72)*.

Consequently, patients suffering from RA and comorbid depression may be highly benefited by the use of Amisulpride. Importantly, clinical use of several dopamine receptor antagonists in schizophrenia patients has been associated with a much lower incidence of RA compared to the general population*(73)*. Accordingly, studying RA patients, who have been under Amisulpride treatment prescribed for their depression symptoms, for possible beneficial clinical and histological RA outcome might provide valuable knowledge for the anti-arthritic effects of the drug. In parallel, preclinical studies of the *in vivo* inhibition of ASCC3 and SEC62 would further corroborate the use of Amisulpride as a repurposing candidate and render it as a promising “lead” compound for the development of novel more potent anti-rheumatic therapeutics.

## 4. Materials and methods

### Amisulpride

For *in vivo* experiments, the commercially available Solian solution (Sanofi, 100 mg/mL) was used. For *in vitro* assays, synthesized Amisulpride was used (100mM stock in DMSO, further dilutions in DMEM)*(74)*.

### SFs isolation

Primary mouse SFs were isolated from ankle joints of 8 weeks CBA;C57BL/6J, WT or *hTNFtg* mice as previously described*(75)*. Briefly, ankle joints were digested with Collagenase IV (Sigma Aldrich, C5138) and at passage 1 a depletion of CD45+ cells was performed using a Biotin anti-mouse CD45 Antibody (Biolegend, 103104) and Dynabeads™ Biotin Binder (ThermoFisher 11047), according to the manufacturer’s protocol.

### Elisa assays

O/N starved *hTNFtg* SFs were treated with different Amisulpride concentrations. WT SFs were stimulated with 10 ng/mL hTNF (Peprotech) or hTNF pre-incubated with Amisulpride for 30 min. 48h after, the cell culture supernatants were analysed for the detection of CCL5 (DY478, DuoSet), CCL20 (DY760, DuoSet), hTNF (R&D Systems, DTA00D) or other chemokines (Mouse 13plex Legendplex Chemokine Panel, Biolegend, 740007) according the the manufacturer’s instructions. The toxicity of Amisulrpide was assessed using the Crystal Violet assay on SFs, as described elsewhere*(76)*.

### L929 Cell Line and TNF induced cell death assay

The L929 cells (NCTC clone, ATCC), TNF–induced cytotoxicity assay was performed as previously described*(77–79)*.

### *In vivo* experiments

All mice were maintained under specific pathogen-free conditions in conventional, temperature-controlled, air-conditioned animal house facilities of BSRC Al. Fleming with 12 h light/12 h dark cycle. The mice received food and water ad libitum. All mice were observed for morbidity and euthanized when needed according to animal welfare.

### LPS model

C57BL/6J mice were challenged with 1μg LPS intraperitoneally. Amisulpride (at 50mg/kg or 100mg/kg in water) was administered to C57BL/6J mice per os (t-2, t0). Sera samples were collected 1.5hrs post induction in order to measure the levels of IL6 and mTNF using the Duoset (DY406) and eBioscience (88-7324) elisas, respectively, following the manufacturer’s protocol.

### *hTNFtg* model

CBA;C57BL/6J, *hTNFtg* mice*(80)* were administered per os 20mg/kg Amisulpride diluted in water, twice daily at 5-8week of age. Arthritis was evaluated clinically in a blinded manner using a semiquantitative arthritis score ranging from 0 to 4, as described elsewhere*(81)*.

### Histology

Formalin-fixed, EDTA-decalcified, paraffin-embedded mouse joint tissue specimens were sectioned and stained with haematoxylin-eosin (H&E), Toluidine Blue (TB) or Tartrate-Resistance Acid Phosphatase (TRAP) Kit (Sigma-Aldrich). H&E and TB were semi-quantitatively blindly evaluated for: synovial inflammation/hyperplasia (scale of 0–5) and cartilage erosion (scale of 0–5) based on an adjusted, previously described method*(82)*. TRAP staining was performed to measure number of osteoclasts using ImageJ software on images acquired with a Nikon microscope, equipped with a QImaging digital camera.

### FACs analysis-Immune infiltration

FACS analysis-based immune infiltration was performed by digesting ankle joints of 8week *hTNFtg* vehicle or Amisulpride treated animals, with 1000U/ml Collagenase IV (Sigma-Aldrich), as previously described*(83)*. 1-2 million cells were stained according to Table 1 of supplementary material.

### RNA sequencing-RNA sequencing analysis

RNA-seq was performed in three biological replicates of cultured SFs treated with 500μM Amisulpride or vehicle for 48h. RNA extraction of cultured SFs was performed using the Qiagen RNA Micro kit (74004), according to the company’s guidelines. RNA extraction of 8w *hTNFtg* mice back joints CD31^-^/ CD45^-^/ PDPN^+^ cells was performed using the Single Cell RNA Purification Kit (Norgen, 51800). All samples were quantified by a Bioanalyzer using the Agilent RNA 6000 Nano Kit reagents and protocol (Agilent Technologies). Only RNA samples with RNA Integrity Number (RIN) >7 proceeded to further analysis. The library preparation was performed according to the 3’ mRNA-Seq Library Prep Kit Protocol for Ion Torrent (QuantSeq-LEXOGEN^™^). The libraries quality was measured in a Bioanalyzer, using the DNA High Sensitivity Kit reagents (Agilent Technologies) according to manufacturer’s protocol, and they were equated at a concentration of 50pM. Templating was performed in the Ion Proton Chef instrument, following the Ion PI™ IC200™ Chef Kit (ThermoFisher Scientific). Sequencing was performed in the Ion Proton™ System according to the Ion PI™ Sequencing 200 V3 Kit on Ion Proton PI™ V2 chips (ThermoFisher Scientific).

For analysis, the raw bam files were summarized to read counts using the Bioconductor package Genomic Ranges. Genes that had zero counts were removed and the gene counts were normalized. Subsequently differential expression analysis was conducted using the Bioconductor package DESeq1*(84)*. Differentially regulated genes were concluded using an absolute log2fold change cutoff value of 1 and a p-value cutoff of 0.05. The aforementioned analytical steps were performed through the Bioconductor package metaseqr*(85)*. R packages*(86)* were used for generating volcano plots, heatmaps, bubble plots and bar plots. Venn diagrams were created with the online available tool InteractiVenn*(87)*. Functional enrichment analysis was done with Enrichr online tool*(88)*.

### TNF/TNFRI Elisa assay

TNF-TNFRI Elisa assay was performed as described elsewhere*(77)*.

### Click compounds synthesis

For “click 1”, the aromatic part of Amisulpride **h** was synthesized*(74)*, which subsequently reacted with 1-Boc-2-(aminomethyl)pyrrolidine **i** under HATU amidation coupling conditions to furnish compound **k**. Removal of the Boc group followed by reaction with propargyl bromide afforded the desired derivative (Figure S4A). Regarding the synthesis of “click 2”, a slightly different procedure was developed including first the synthesis of the iodide **n**, which upon palladium-catalysed multicomponent reductive cross coupling reaction with 4-bromo-1-butyne and sodium metabisulfite as an inorganic sulfur dioxide surrogate*(89)* afforded compound **o**. Hydrolysis of the ester derivative followed by amidation coupling with 2-(aminomethyl)-1-ethylpyrrolidine yielded finally the “click 2” derivative (Figure S4B).

### Mass spectrometry experiments for target identification

In order to identify the possible targets of Amisulpride on *hTNFtg* SFs a previously described method*(90)* was employed, by lowering down the amount of total protein used at 4mg. A brief description of the protocol followed is provided in the supplementary material and methods. The samples digestion, run and initial analysis were processed as previously described*(91)*. A summary of the process followed is described in the supplementary material and methods.

### shRNAS and Lenti-X transfection

For silencing of potential targets of Amisulpride, *hTNFtg* SFs were transduced by Lenti viruses expressing a full list of shRNAs (described in Supplementary table 2). ShRNAs were cloned in the HpaI/ XhoI sites of PLB vector. To produce the viruses HEK Lenti-X 293T cell line (Clontech) were transfected using PEI (Sigma, P-3143). Plasmids pLB, psPAX2, pMD2G were purchased from Addgene. The lentiviral transduction of SFs was performed using Hexadimethrine Bromide/ Polybrene (Sigma, H9268) and transduction efficiency was calculated by the FITC positive cells in FACs Canto II Flow cytometer (BD Biosciences) and FlowJo software (FlowJo, LLC). Sorting of the GFP positive cells was performed, when transfection efficiency was less than 70%. SFs tranduced by lentiviruses that carry a vector containing scramble shRNA, were used as relevant control (Supplementary table 2).

### Gene expression quantification

To confirm several interesting candidates of RNAseq results, to test if known targets of Amisulrpide are expressed in SFs or to investigate if tranduced SFs underexpress the gene of interest, we performed a qPCR analysis using the Platinum SYBR-Green qPCR SuperMix (Invitrogen), after CDNA synthesis by the MMLV Reverse Transcriptase (Promega). The CFX96 Touch Real-Time PCR Detection System (Biorad) was used and quantification was performed with the DDCt method. Primer sequences (5′-3′) are provided in Table 3 of Supplementary Material.

### Phosphoproteomics samples preparation and analysis

WT SFs activated by hTNF for 5’, 15’ 30’ with or without pre-incubation with 500μM Amisulpride, 1h before in triplicates were processed as previously described*(92)*. Further process of samples and subsequent analysis is described in supplementary material.

### Statistical analysis

All experiments were performed at least 3 times. Data are presented as mean ± SE. Student’s *t*-test (parametric, unpaired, two-sided) or two-way ANOVA were used for evaluation of statistical significance using GraphPad Prism 8 software. Statistical significance is presented as follows: * p < 0.05, ** p < 0.01, *** p < 0.001.

## Ethics declarations

All experiments were approved by the Institutional Committee of Protocol Evaluation in conjunction with the Veterinary Service Management of the Hellenic Republic Prefecture of Attica according to all current European and national legislation. The authors declare no competing financial interests.

## Contributions

All authors read the manuscript and provided feedback. D.P. conducted the experiments, collected the data and prepared the manuscript. F.R. performed the clinical arthritis scoring and contributed in FACs experiments and Lentiviruses experiments. C.T. and P.C. performed the bioinformatics analysis. A.K.P. performed the phosphoproteomics experiments. F.C., E.C.V. and L.N. contributed in the *in vivo* experiments. N. K. and M.D. supervised the *in vivo* experiments. J.V.O. supervised the phosphoproetomics studies. A.N.M. designed, analysed and supervised the chemoproteomics experiments and synthesized the two “click” compounds and Amisulpride used in the *in vitro* assays. G.K. designed the study, edited the manuscript and supervised the study.

## Acknowledgements

The authors thank P. Athanasakis for breeding the mice and A. Katevaini for excellent technical assistance in the histology part of the paper. Authors also thank M. Samiotaki and G. Stamatakis for generating and analysing the chemoproteomics raw data. Authors also appreciate the contribution of S. Grammenoudi and K. Dagla in running the FACs sorting experiments and V. Harokopos for Genomics service.

The authors also acknowledge support of this work by projects Drug.Art (T2EΔK-01076) and InfrafrontierGR/Phenotypos (MIS 5002135), funded by the Operational Programme “Competitiveness, Entrepreneurship and Innovation” (NSRF 2014-2020) co-financed by Greece and the European Union (European Regional Development Fund). Finally, this work has been supported by EPIC-XS, project number 0001017, funded by the Horizon 2020 programme of the European Union.

