## Supplementary Figures and Methods for "Repurposing of Amisulpride, a known antipsychotic drug, to target synovial fibroblasts activation in arthritis"

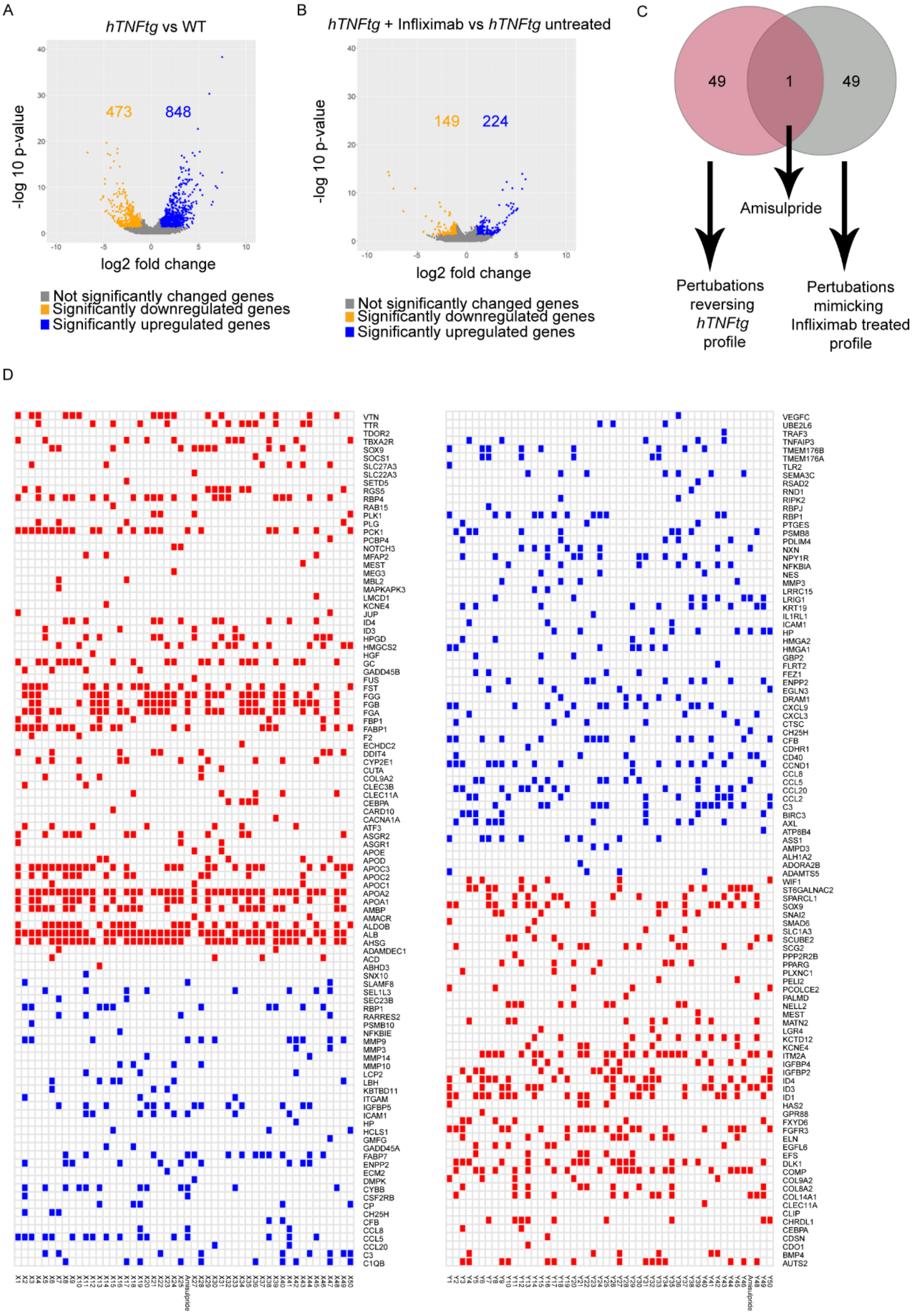


**Figure S1 Methodology for the identification of new candidates for the treatment of RA** A) Volcano plot showing the number of deregulated genes in *hTNFtg* SFs versus WT control. B) Volcano plot showing the number of deregulated genes in *hTNFtg* SFs before and after treatment with Infliximab. C) Venn diagram showing the overlap between perturbations that reverse the *hTNFtg* profile and mimic the Infliximab-treated disease signature. D) Heatmaps presenting the top 50 perturbations proposed by the L1000CDS$^2$ search engine, that either reverse the *hTNFtg* profile or mimic the Infliximab-treated disease signature, respectively.

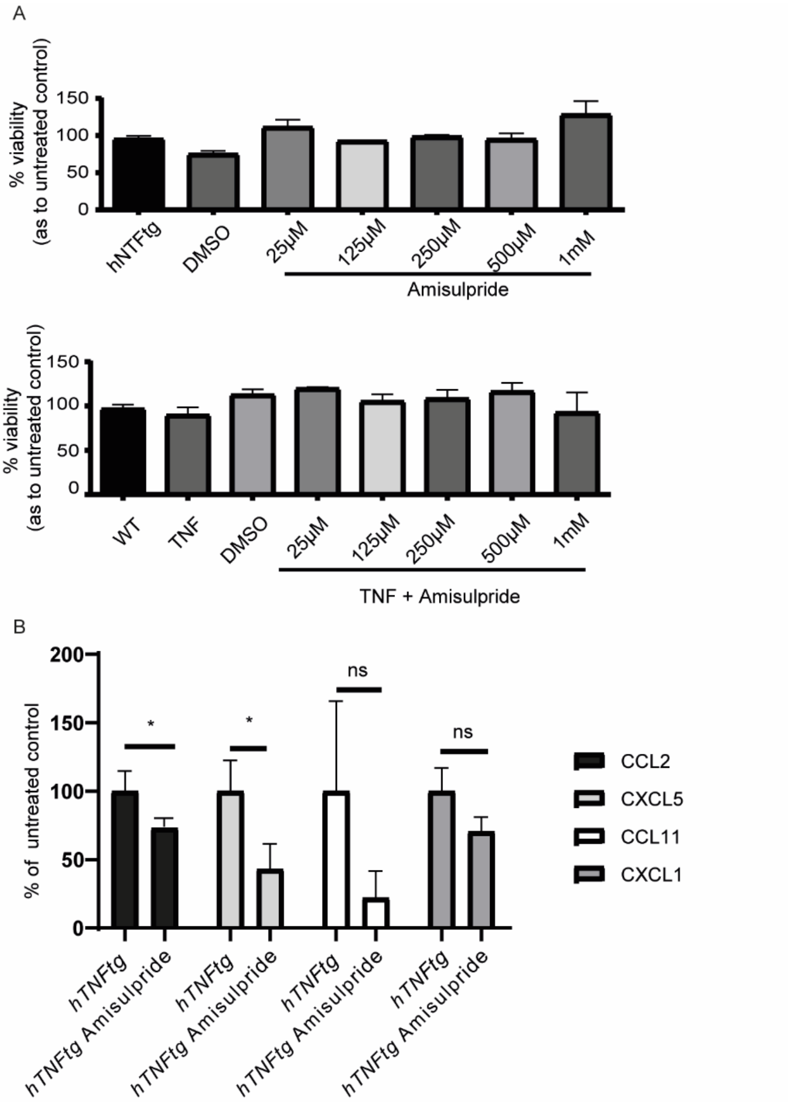


**Figure S2** A. Crystal violet viability assay, indicating the non-toxicity of Amisulpride, even in high doses. B. Amisulpride at 500µM downregulates CCL2 (MCP1) and CXCL5 (LIX), measured by Legendplex panel on *hTNFtg* SFs and TNF stimulated WT SFs. CCL11 and CXCL1 were not downregulated significantly. TARC (CCL17), MIP-1β (CCL4), BLC (CXCL13) and MDC (CCL22) were not detected in the supernatants. (* p-value < 0.05; ** p-value < 0.01; *** p-value ≤ 0.0001, all data are shown as mean ± SEM and all comparisons were made against *hTNFtg* vehicle sample using Student's t test.)

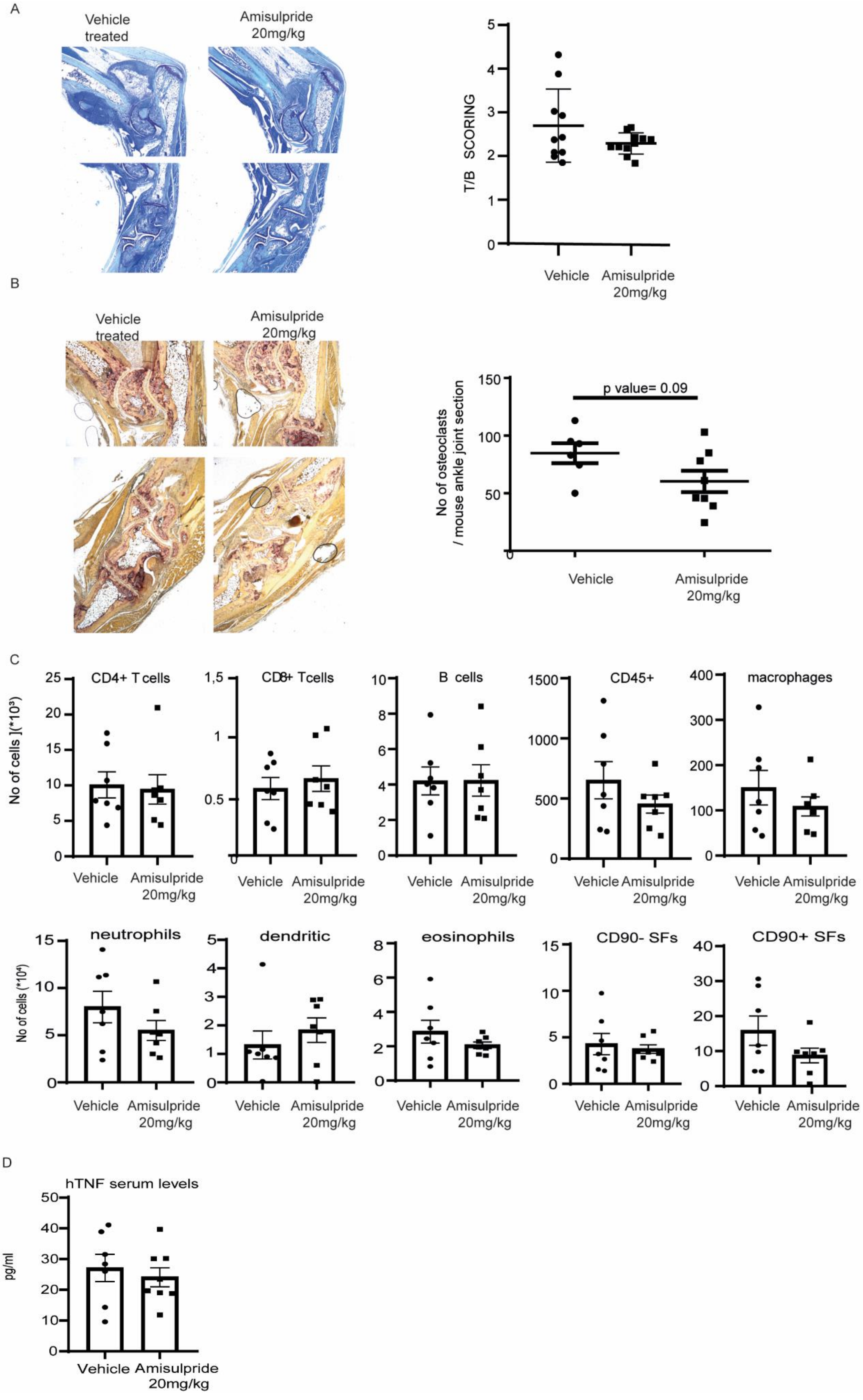


**Figure S3** *In vivo* A. T/B and B. TRAP staining of joints paraffin sections show a non-significant effect of Amisulpride treatment in *hTNFtg* animals on cartilage destruction and bone erosion, respectively. C. Immune infiltration FACs analysis of joints of *hTNFtg* mice treated with Amisulpride for 5weeks, compared with the vehicle treated control D. hTNF serum do not differ in mice treated with Amisulpride (* p-value < 0.05; ** p-value < 0.01; *** p-value ≤ 0.0001, all data are shown as mean ± SEM and all comparisons were made against *hTNFtg* vehicle sample using Student's t test.)

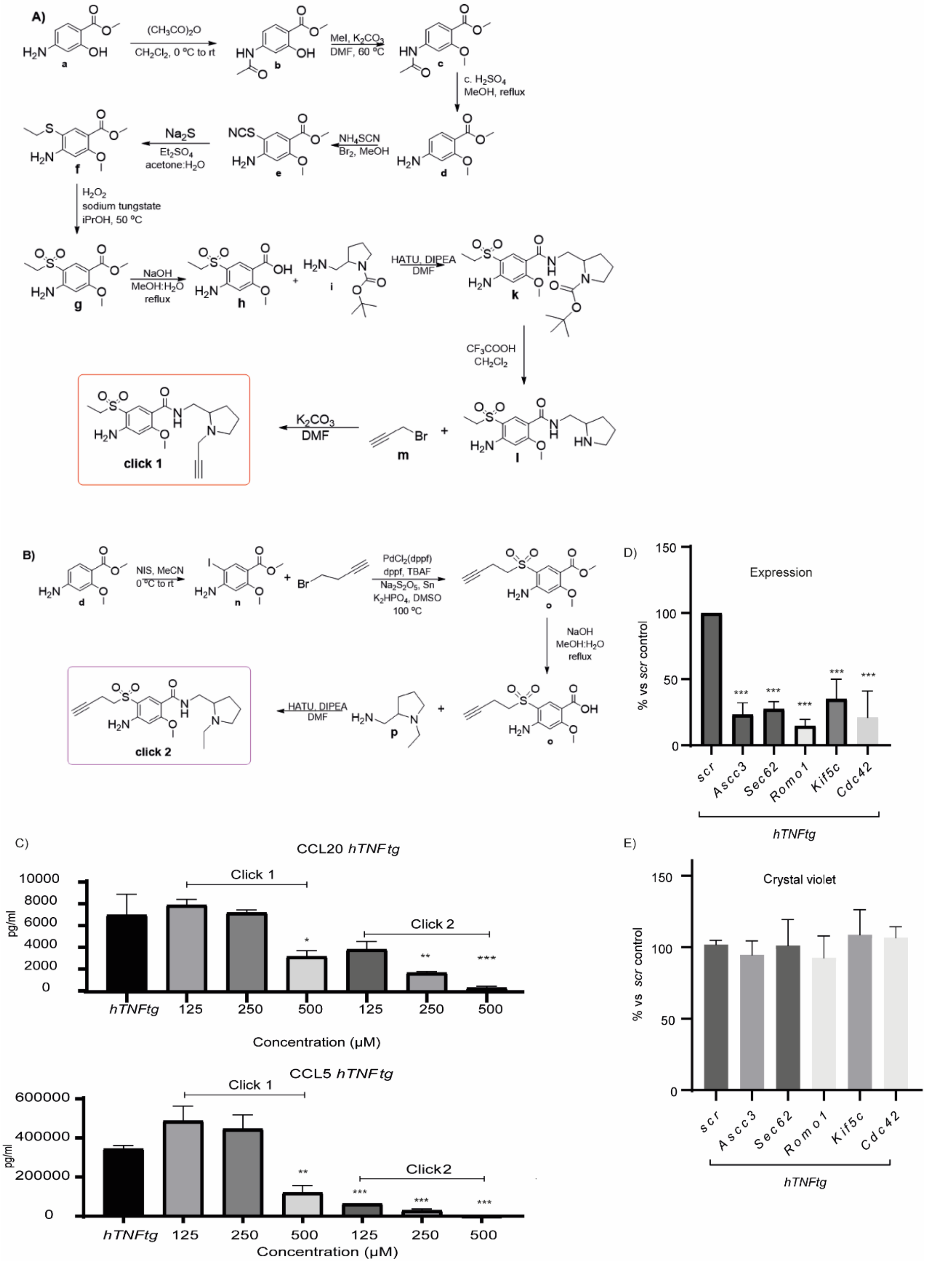


**Figure S4**. Synthetic routes followed for the synthesis of compounds A. click 1 and B. click 2. C. Click compounds maintain their efficacy in downregulating CCL5 and CCL20 at *hTNFtg* arthritic SFs. D. Efficient downregulation of Amisulpride potential targets, upon Lentiviral transfection of *hTNFtg* SFs with relevant shRNAs. E. Crystal violet assay confirms that the shRNAs transfection is not toxic. (* p-value < 0.05; ** p-value < 0.01; *** p-value ≤ 0.0001, all data are shown as mean ± SEM and all comparisons were made against *hTNFtg* vehicle sample using Student's t test.)

### NMR spectra

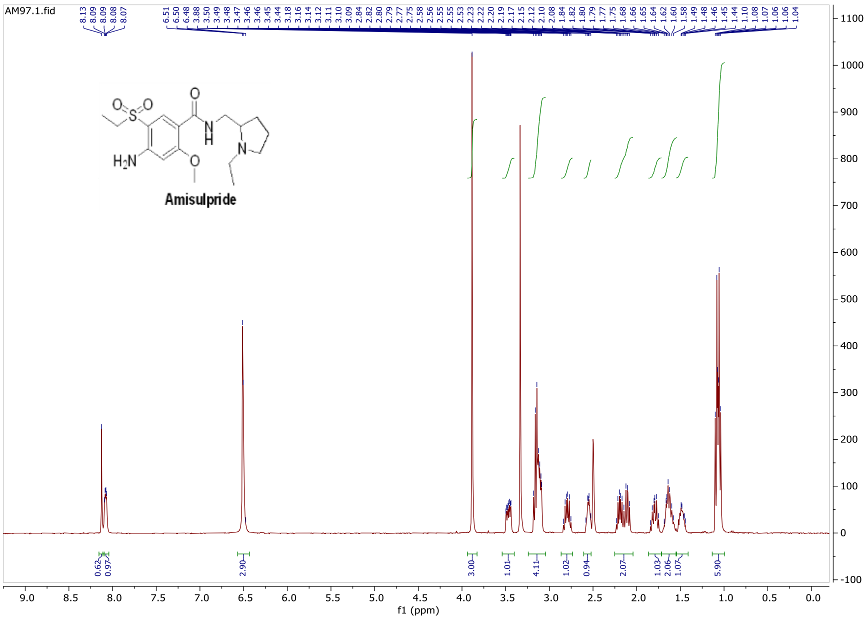


$^{1}H$-NMR (400 MHz, dmso-$d_6$) $\delta$ 1.07 (dt, $J_1$ = 10.6 Hz, $J_2$ = 7.2 Hz, 6H), 1.44-1.49 (m, 1H), 1.58-1.68 (m, 2H), 1.75-1.84 (m, 1H), 2.08-2.23 (m, 2H), 2.53-2.58 (m, 1H), 2.75-2.84 (m, 1H), 3.09-3.18 (m, 4H), 3.47 (ddd, $J_1$ = 13.4 Hz, $J_2$ = 6.9 Hz, $J_3$ = 2.9 Hz, 1H), 3.88 (s, 3H), 6.48-6.51 (m, 3H), 8.07-8.09 (m, 1H), 8.13 (s, 1H). MS [ESI$^+$] m/z: 370.5 [M + H$^+$]$^+$.

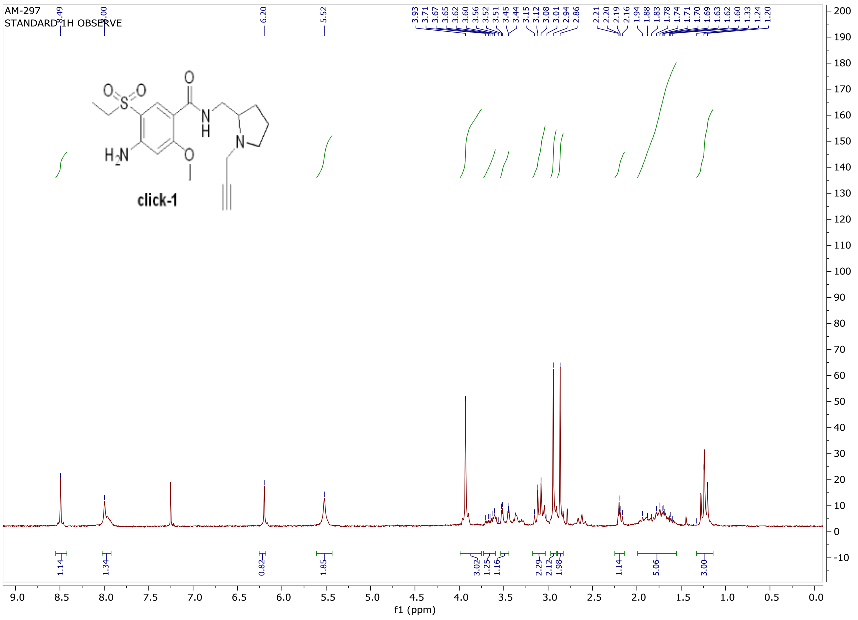


$^{1}H$-NMR (200 MHz, $CDCl_3$) $\delta$ 1.24 (t, $J$ = 7.2 Hz, 3H), 1.60-1.94 (m, 5H), 2.16-2.21 (m, 1H), 2.86 (s, 2H), 2.94 (s, 2H), 3.01-3.15 (m, 2H), 3.44-3.52 (m, 1H), 3.56-3.71 (m, 1H), 3.93 (s, 3H), 5.52 (brs, 2H), 6.20 (s, 1H), 8.00 (brs, 1H), 8.49 (s, 1H). MS [ESI$^+$] m/z: 380.6 [M + H$^+$]$^+$.

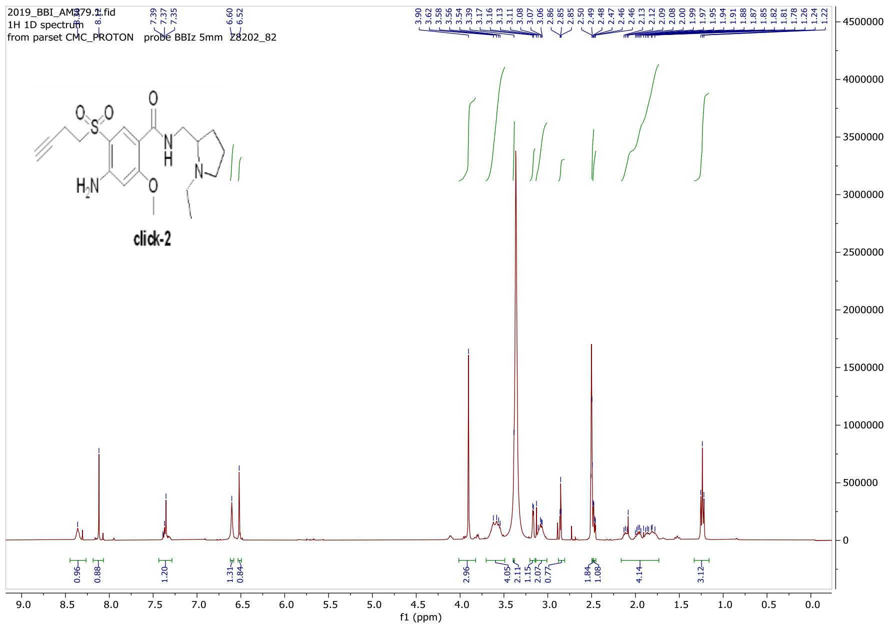


$^1$H-NMR (400 MHz, dmso-d$_6$) $\delta$ 1.24 (t, $J$ = 7.2 Hz, 3H), 1.78-2.13 (m, 4H), 2.47 (dd, $J_1$ = 7.4 Hz, $J_2$ = 2.8 Hz, 1H), 2.48-2.50 (m, 2H), 2.85 (t, $J$ = 2.7 Hz, 1H), 3.06-3.13 (m, 2H), 3.17 (d, $J$ = 3.6 Hz, 1H), 3.39 (m, 2H), 3.54-3.62 (m, 4 H), 3.90 (s, 3H), 6.52 (s, 1H), 6.60 (s, 1H), 7.35-7.39 (m, 1H), 8.12 (s, 1H), 8.30 (s, 1H). MS [ESI$^+$] m/z: 394.18 [M + H$^+$]$^+$.

**Figure S5.** $^1$H-NMR spectra and relevant analysis for Amisulpride and its click derivatives (click-1 and click-2) which were used in the chemoproteomic analysis.

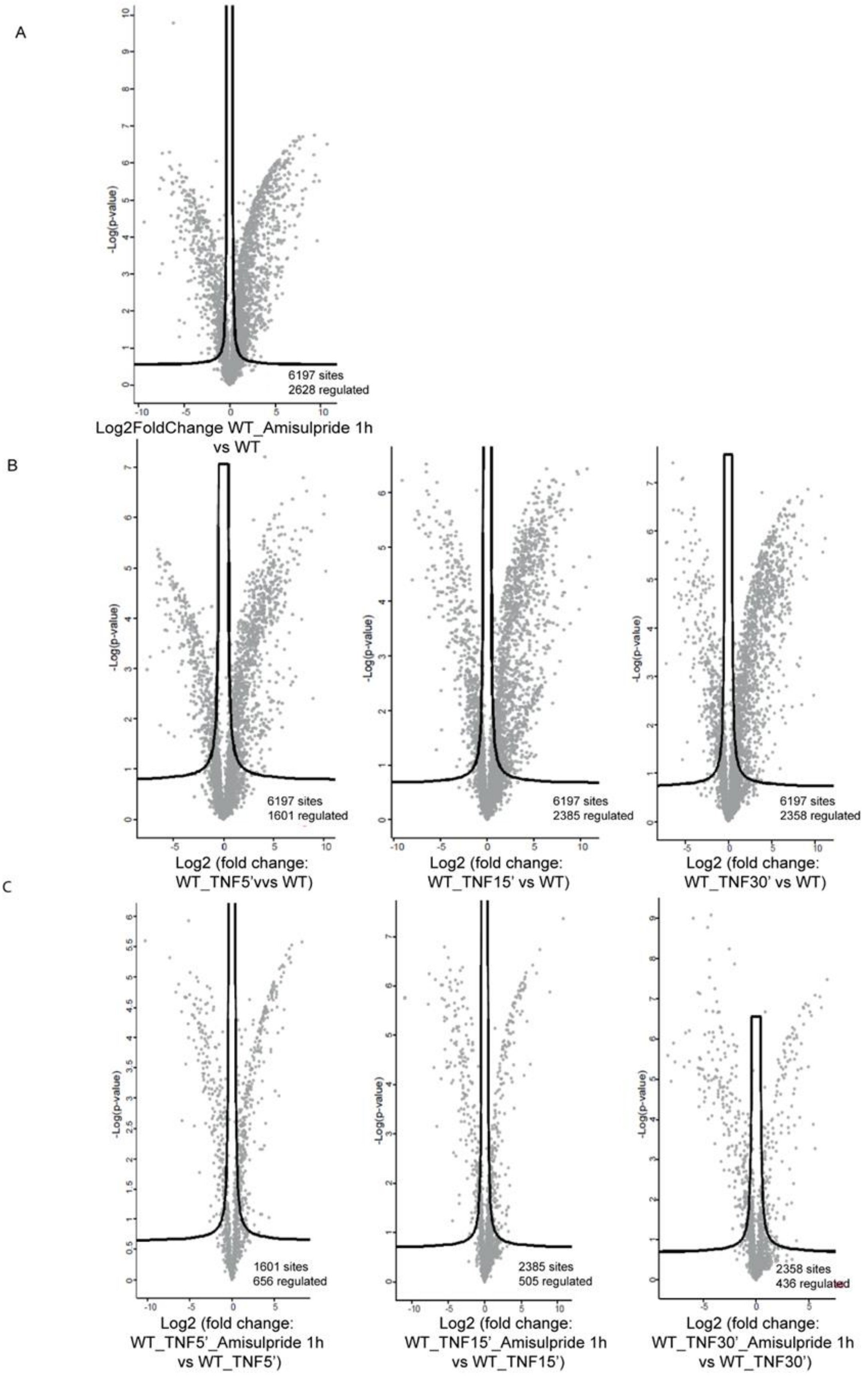


**Figure S6** A. Volcano plot illustrating phosphorylations that are up and down regulated upon Amisulpride treatment with significance cut-off = 0.05 indicated by lines. B. Volcano plots indicating the deregulation of phosphoproteome on hTNF stimulated WT SFs, when treated with Amisulpride. The samples were gathered on the three indicated timepoints of 5’, 15’ and 30’ after hTNF stimulation and were compared to untreated control. C. Regulated proteins by TNF treatment, identified in B, were used as input to compare corresponding inhibitor treatment.

### Supplementary Materials and methods

#### Antibodies used in Immune infiltration FACs analysis

Unspecific binding was blocked by the anti-Fc Receptor (anti-CD16/32) antibody (Biolegend). Analysis was performed using a FACS Canto II Flow cytometer (BD Biosciences) and FlowJo software (FlowJo, LLC). Counting beads were used for quantification of different cell subsets. Supplementary Table 1 summarizes the antibodies used for FACs staining.

| **Cells subset** | **Antibodies (Company, Cat. Number)** | | | | | | | | **Live/ Dead Exclusion** |
|---|---|---|---|---|---|---|---|---|---|
| **Myeloid cells** | PE-conjugated anti-CD11b (BD Biosciences, 557397) | A700-conjugated anti-CD45 (Biolegend, 103128) | APC-conjugated anti-MHCII (eBioscience,17-5320-82) | PE/Dazzle594-conjugated anti-CD64 (Biolegend, 139320) | APCFire-conjugated anti-CD24 (Biolegend,101840) | PE/Cy7-conjugated anti-CD11c (Biolegend, 117318) | FITC-conjugated anti-Ly6C (BD Biosciences, 553104) | Biotinylated anti-Ly6G (eBioscience, 13-5931-75) with streptavidin-conjugated PE/Cy5 (Invitrogen) | Dapi (Invitrogen, D1306) |
| **Lymphocytes** | PE-conjugated anti-B220 (BD Biosciences, 553089) | ApcCy7-conjugated anti-CD45 (Biolegend, 103116) | PE/Cy7-conjugated anti-CD3 (eBioscience,25-0031-82) | A700-conjugated anti-CD4 (Biolegend, 100536) | APC-conjugated anti-CD8 (Biolegend,100711) | | | | Zombie Green (Sigma, 423112) |
| **Fibroblasts** | A488-conjugated anti-CD90.2 (Biolegend, 105316) | A700-conjugated anti-CD45 (Biolegend, 103128) | PE-conjugated anti-CD31 (BD Biosciences, 553373) | PE/Cy7-conjugated anti-PDPN (Biolegend, 127412) | | | | | Zombie NIR (Sigma, 77184) |

**Supplementary Table 1:** Antibodies used in Immune infiltration FACs analysis

### ShRNAs sequences for Lentiviral vectors creation

| Gene to silence | shRNA sequence |
|---|---|
| ***Ascc3*** | 5' T-GGAAGAAGATAGTGAAATT-TTCAAGAGA-AATTTCACTATCTTCTTCC-TTTTTTC 3' |
| | 3' A-CCTTCTTCTATCACTTTAA-AAGTTCTCT-TTAAAGTGATAGAAGAAGG-AAAAAAGAGCT 5' |
| ***Kif5c*** | 5' T-GCAAAGACCATCAAGAATA-TTCAAGAGA-TATTCTTGATGGTCTTTGC-TTTTTTC 3' |
| | 3' A-CGTTTCTGGTAGTTCTTAT-AAGTTCTCT-ATAAGAACTACCAGAAACG-AAAAAAGAGCT 5' |
| ***Romo1*** | 5' T-GCAAAGACCATCAAGAATA-TTCAAGAGA-TATTCTTGATGGTCTTTGC-TTTTTTC 3' |
| | 3'A-CCCTTTTGGTACTACGTCT-AAGTTCTCT-AGACGTAGTACCAAAAGGG-AAAAAAGAGCT 5' |
| ***Sec62*** | 5' T-GGGATTAATTCTTGTGATT-TTCAAGAGA-AATCACAAGAATTAATCCCT-TTTTTC 3' |
| | 3' A-CCCTAATTAAGAACACTAA-AAGTTCTCT-TTAGTGTTCTTAATTAGGG-AAAAAAGAGCT 5' |
| ***Cdc42*** | 5' T-GCAAGAGGATTATGACAGA-TTCAAGAGA-TCTGTCATAATCCTCTTGC-TTTTTTC 3' |
| | 3' A-CGTTCTCCTAATACTGTCT-AAGTTCTCT- AGACAGTATTAGGAGAACG-AAAAAAGAGCT 5' |

**Supplementary Table 2:** shRNAs sequences used in the study.

### Real time Primers' sequences

The table below describes the 5' to 3' sequences of RT primers used. B2m or Gapdh were used as reference genes.

| Gene | 5'→ 3'sequence |
|---|---|
| *hTNF* | 5'-CTTCTCGAACCCCGAGTGAC-3' |
| *Mmp3* | 5'-GTCTCCCTGCAACCGTGAA-3' |
| *Cxcl3* | 5'-CTGCACCCAGACAGAAGTCATA-3' |
| *Cox2* | 5'-TCAGTTTTTCAAGACAGATC-3' |
| *DRD2* | 5'-GTTTCCCAGTGAACAGGCGG-3' |
| *DRD3* | 5'-TGGCAACGGTCTGGTATGTG -3' |
| *Htr7* | 5'-TGCAACGTCTTCATCGCCA-3' |
| *Ascc3* | 5'-GGCCTTACATGGAAGAAGATAGTG-3' |
| *Kif5c* | 5'-AGCGGGAAGCTGTATTTGGT-3' |
| *Romo1* | 5'-GAGCACTCTCGCCGCAGAT-3' |
| *Sec62* | 5'-GCAGTAATAGCAGCCACCCT-3' |
| *Cdc42* | 5'-GGCGGAGAAGCTGAGGAC-3' |
| *B2m* | 5'-TTCTGGTGCTTGTCTCACTGA-3' |
| *Gapdh* | 5'-GTTGTCTCCTGCGACTTCA-3' |

**Supplementary Table 3:** 5' to 3' sequences of RT primers used in the study.

**Mass spectrometry experiments for target identification - SP3 digestion and clean up- LC-MS/MS- Data analysis**

In brief, after cell treatment and cell lysis, the lysate was subjected to the copper(I)-catalysed alkyne-azide cycloaddition click chemistry approach, for conjugating target proteins with a biotin tag. The biotin-tagged proteins were then pulled down on streptavidin beads, and the target proteins were selectively eluted through cleaving an azo-linker in the tag with sodium dithionite. Finally, the proteins enriched in the eluent were identified by mass spectrometry analysis.

In summary, the eluates were processed using the sensitive sp3 protocol*(1)*, peptide mixtures were collected and were loaded on the trap column at 10uL/min for 4 min with 0.1% formic acid in water and separated in a gradient of 0.1% (vol/vol) formic acid in mobile phase A. and B. acetonitrile.

The data acquisition was performed in positive mode using a Q Exactive HF-X Orbitrap mass spectrometer (ThermoFisher Scientific). MS data were acquired in a data-dependent strategy and the resolution of the survey scan was 120,000 (at m/z 200) with a target value of 3 × 10E6 ions and a maximum injection time of 100 ms.

The acquired raw files were processed through the MaxQuant software (1.6.14.0) using the Mus musculus proteome FASTA database. Perseus (version 1.6.10.43) was used and proteins identified as 'contaminants', 'reverse' and 'only identified by site' were filtered out. The three replicates of each condition were grouped (treated versus vehicle/ competition condition).  A two-sided Student's t-test of the grouped proteins was performed using p value <0.05 as a significance measurement.

**Phosphoproteomics samples processing and analysis**

Cell pellets were lysed in buffer containing 5% sodium dodecyl sulfate (SDS), 5 mM tris(2-carboxyethyl)phosphine (TCEP), 10 mM chloroacetamide (CAA), 100 mM Tris, pH 8.5 and boiled for 10' followed by sonication with a micro tip probe. Protein concentrations were estimated using the Pierce BCA Protein Assay Kit (ThermoFisher Scientific). 500 µg of protein/ sample was used for PAC digestion in an automated 96-well format on a KingFisher™

Flex robot (Thermo Fisher Scientific) with 12 hour O/N digestion at 37 °C, using LysC (Wako) and Trypsin (Sigma Aldrich) as described before*(2)*. Protease activity was quenched by acidification with trifluoroacetic acid (TFA). Peptide mixtures were purified and concentrated on reversed-phase C18 Sep-Pak cartridges (Waters).

For phosphoproteome analysis enrichment of phosphopeptides was carried out in 96-well format on a KingFisher™ Flex robot (Thermo Fisher Scientific) based on previously described protocols*(2, 3)*. Peptides were eluted with 75 µl of 80% acetonitrile directly into a KingFisher 96-well plate and subsequently with 150 µl of loading buffer (80% ACN, 8% TFA and 1.6 M glycolic acid). Phosphopeptides were enriched using TiIMAC-HP beads (MagResyn, Resyn Biosciences) and eluted in the final plate with 1% ammonia. After the eluted phosphopeptides were acidified with TFA, they were directly loaded onto EvoTips according to the manufacturer's protocol.

All samples were analyzed on the EvoSep One system (using the pre-programmed 60 samples/ day gradient) coupled to an Orbitrap Exploris 480 MS (Thermo Fisher Scientific) through a nanoelectrospray source. Peptides were separated on a 15-cm, 150 µM inner diameter analytical column in-house packed with 1.9 µM reversed-phase C18 beads (ReProsil-Pur AQ, Dr Maisch) and column temperature was maintained at 60°C by an integrated column oven (PRSO-V1, Sonation GmbH). Phosphoproteome analysis was performed using data-independent acquisition (DIA).

All DIA raw files were analyzed using Spectronaut with a library-free approach (directDIA). All files were searched against the mouse UniProt database, supplemented with commonly observed contaminants. For phosphoproteome analysis phosphorylation of serine, threonine and tyrosine were included as variable modifications and PTM localization cutoff was set to 0.75. Phospho-peptide data was collapsed to site information using the Perseus plugin previously described*(4)*.

DIA phosphoproteome data were processed using R (version 3.6.2) with the Prostar data analysis pipeline*(5)*. Data were log2 transformed and filtered (a minimum of two valid values in at least one condition were required for an identification to be included in downstream analysis). Data were normalized by quantile-based normalization and imputation to replace missing values was performed using a two-step approach *(3)*. Further data analysis of proteomics data was performed using Perseus software version 1.6.2.2 or 1.6.5.0. Data were normalized by row-based median subtraction and heatmaps were generated based on unsupervised hierarchical clustering. Specifically, to assess overall Amisulpride effect, phosphoproteome data were median normalized within groups defining initial treatment (WT, TNF treated) and significantly regulated phosphosites comparing inhibitor (treated and untreated) were identified by t-test using a significance cut-off of 0.05. Volcano plots were generated for visualization of significantly regulated sites identified by Student's t-test (significance cut-off 0.05). Gene ontology (GO) term enrichment analysis and Kyoto Encyclopedia of Genes and Genomes (KEGG) pathway enrichment analysis were performed using DAVID*(6, 7)* and InnateDb*(8)*, respectively.

**References (Supplementary Material)**


1. C. S. Hughes, S. Foehr, D. A. Garfield, E. E. Furlong, L. M. Steinmetz, J. Krijgsveld, Ultrasensitive proteome analysis using paramagnetic bead technology. *Mol. Syst. Biol.* (2014), doi:10.15252/msb.20145625.

2. D. B. Bekker-Jensen, A. Martínez-Val, S. Steigerwald, P. Rüther, K. L. Fort, T. N. Arrey, A. Harder, A. Makarov, J. V. Olsen, A compact quadrupole-orbitrap mass spectrometer with FAIMS interface improves proteome coverage in short LC gradients. *Mol. Cell. Proteomics* (2020), doi:10.1074/mcp.TIR119.001906.

3. A. Martinez-Val, D. B. Bekker-Jensen, S. Steigerwald, C. Koenig, O. Østergaard, A. Mehta, T. Tran, K. Sikorski, E. Torres-Vega, E. Kwasniewicz, S. H. Brynjólfsdóttir, L. B. Frankel, R. Kjøbsted, N. Krogh, A. Lundby, S. Bekker-Jensen, F. Lund-Johansen, J. V. Olsen, Spatial-proteomics reveals phospho-signaling dynamics at subcellular resolution. *Nat. Commun. 2021 121* (2021).

4. D. B. Bekker-Jensen, O. M. Bernhardt, A. Hogrebe, A. Martinez-Val, L. Verbeke, T. Gandhi, C. D. Kelstrup, L. Reiter, J. V. Olsen, Rapid and site-specific deep phosphoproteome profiling by data-independent acquisition without the need for spectral libraries. *Nat. Commun.* (2020), doi:10.1038/s41467-020-14609-1.

5. S. Wieczorek, F. Combes, C. Lazar, Q. G. Gianetto, L. Gatto, A. Dorffer, A. M. Hesse, Y. Couté, M. Ferro, C. Bruley, T. Burger, DAPAR & ProStaR: Software to perform statistical analyses in quantitative discovery proteomics. *Bioinformatics* (2017), doi:10.1093/bioinformatics/btw580.

6. D. W. Huang, B. T. Sherman, R. A. Lempicki, Systematic and integrative analysis of large gene lists using DAVID bioinformatics resources. *Nat. Protoc.* (2009), doi:10.1038/nprot.2008.211.

7. D. W. Huang, B. T. Sherman, R. A. Lempicki, Bioinformatics enrichment tools: Paths toward the comprehensive functional analysis of large gene lists. *Nucleic Acids Res.* (2009), doi:10.1093/nar/gkn923.

8. D. J. Lynn, G. L. Winsor, C. Chan, N. Richard, M. R. Laird, A. Barsky, J. L. Gardy, F. M.

Roche, T. H. W. Chan, N. Shah, R. Lo, M. Naseer, J. Que, M. Yau, M. Acab, D. Tulpan, M. D. Whiteside, A. Chikatamarla, B. Mah, T. Munzner, K. Hokamp, R. E. W. Hancock, F. S. L. Brinkman, InnateDB: Facilitating systems-level analyses of the mammalian innate immune response. *Mol. Syst. Biol.* (2008), doi:10.1038/msb.2008.55.
